# Comprehensive genome editing confers ‘off-the-shelf’ CAR-T cells superior efficacy against solid tumors

**DOI:** 10.1101/2023.08.03.551705

**Authors:** Ryan Murray, Nuria Roxana Romero Navarrete, Kashvi Desai, Md Raihan Chowdhury, Shanmuga Reddy Chilakapati, Brian Chong, Angelica Messana, Hanna Sobon, Joe Rocha, Faith Musenge, Adam Camblin, Giuseppe Ciaramella, Michail Sitkovsky, Colby Maldini, Stephen Hatfield

## Abstract

Biochemical and immunological negative regulators converge to inhibit tumor-reactive Chimeric Antigen Receptor T (CAR-T) cells, which may explain clinical failures of CAR-T cell therapies against solid tumors. Here, we developed a multifaceted approach to genetically engineer allogeneic (‘off -the-shelf’) CAR-T cells resistant to both biochemical (adenosine) and immunological (PD-L1 and TGF-β) inhibitory signaling. We multiplexed an adenine base editor with a CRISPR-Cas12b nuclease to manufacture a CAR-T cell product comprising six gene edits to evade allorejection (*B2M, CIITA*), prevent graft-versus-host disease (*CD3E*) and resist major biochemical (*ADORA2A*) and immunological (*PDCD1*, *TGFBR2*) immunosuppressive barriers in solid tumors. Combinatorial genetic disruption in CAR-T cells enabled superior anti-tumor efficacy leading to improved tumor elimination and survival in humanized mouse models that recapitulated the suppressive features of a human tumor microenvironment (TME). This novel engineering strategy conferred CAR-T cells resistance to a diverse TME, which may unlock the therapeutic potential of CAR-T cells against solid tumors.

**One Sentence Summary:** Multiplex genome engineered CAR-T cells resistant to allorejection and the convergence of biochemical and immunological negative regulators within the tumor microenvironment exhibit superior efficacy against solid tumors.

## Introduction

Chimeric Antigen Receptor T (CAR-T) cell therapies have elicited remarkable clinical responses in treating hematological malignancies [1,2]. However, their effectiveness in treating solid tumors remains limited due to the complex immunosuppressive tumor microenvironment (TME) [3,4]. Tumor hypoxia, resulting from rapid tumor growth and irregular vasculature [5–8] stabilizes hypoxia-inducible factor 1-alpha (HIF-1α) and promotes the generation of an adenosine-rich TME [9–12]. Extracellular adenosine activates adenosine A_2A_ receptors (A_2A_R) expressed on T cells and potently suppresses T cell effector functions [13–18]. Alongside adenosine-mediated biochemical suppression, the heterogeneous organization of the TME presents immunological barriers impeding CAR-T cell efficacy. For instance, hypoxia/HIF-1α increases the expression of PD-L1 and TGF-β [19,20] that signal through PD-1 and TGF-βRI/RII, respectively, to attenuate T cell activity [21,22]. Prior attempts to improve CAR-T cell efficacy by disrupting single genes involved in these pathways have demonstrated promising preclinical efficacy [23–30].

However, the development of a CAR-T cell product overcoming multiple inhibitory pathways may have distinct advantages in responding to intra-/inter-patient tumor heterogeneity. Here, we leverage base editing to address current challenges in developing multiplex gene-edited CAR-T cell products. Base editors (BE) have enabled precise base pair changes to disrupt gene expression without inducing double-strand DNA breaks or karyotypic abnormalities [31–33] characteristic of current CRISPR nuclease systems [34,35]. While BEs have been utilized to generate clinical -stage CD7-directed CAR-T cells (NCT05885464) [36], the number of accessible BE target sites containing neighboring NGG protospacer adjacent motifs (PAMs) may be restricted for any given gene. Thus, to broaden the range of targeting loci and develop a higher-order gene-edited CAR-T cell product, it may be advantageous to combine base editing with a CRISPR nuclease. To this end, we complexed our adenine BE (ABE) with a Cas12b nuclease [37], allowing for the elimination of biochemical, immunological and allogeneic barriers resulting in a 6-plex gene-edited solid tumor CAR-T cell product, termed Stealth-TKO CAR-T cells.

Stealth-TKO CAR-T cells may offer clinical promise as they are i) *‘off-the-shelf’* by overcoming challenges associated with autologous CAR-T cell manufacturing and patient accessibility, ii) *safe* by avoiding graft-versus-host disease (GvHD), and iii) *effective* against solid tumors due to the genetic ablation of critical T cell inhibitory pathways. Stealth-TKO CAR-T cells are resistant to adenosine (A_2A_R-KO), PD-L1 (PD-1-KO) and TGF-β (TGFBR2-KO), which addresses the convergence of biochemical and immunological negative regulators in the TME. Furthermore, we introduced three additional ‘Stealth’ base edits to generate allogeneic CAR-T cells that avoided GvHD and evaded host immunologic rejection by removing the T cell receptor (CD3E-KO), and HLA class-I (B2M-KO) and HLA class-II (CIITA-KO), respectively. Cumulatively, Stealth-TKO CAR-T cells exhibited superior anti-tumor efficacy in stringent humanized xenograft murine systems, underscoring the power of multiplex gene editing to putatively overcome TME-associated immunosuppressive pathways.

## Results

### Adenosine-resistant CAR-T cells demonstrate improved effector function *in vitro*

The accumulation of extracellular adenosine (Ado) in the solid tumor microenvironment (TME) is a powerful biochemical barrier inhibiting effective anti-tumor T cell responses [38–40]. Therefore, we engineered Ado-resistant CAR-T cells by ablating surface expression of the adenosine A_2A_ receptor (A_2A_R) encoded by *ADORA2A*. Single guide RNAs (sgRNAs) were designed to home adenine (ABE) and cytosine (CBE) base editors to *ADORA2A* and alter protein expression by mutating either the start codon, conserved intron-exon mRNA splice motifs, or by installing premature termination codons (**Fig. 1a**). To identify an sgRNA and base editor combination that mediates optimal *ADORA2A* editing, mRNAs encoding ABE or CBE were paired with a corresponding sgRNA and electroporated into activated primary human T cells (**Fig. 1b; Supplementary Table 1**). This screen identified an ABE-sgRNA (TSBTx2043) complex targeting a splice intron-exon junction that achieved a mean on-target genomic editing efficiency of 88% (**Fig. 1c**). Next, we determined whether *ADORA2A* disruption prevents A_2A_R signaling by generating EGFR-specific, 4-1BB co-stimulated CAR-T cells containing the *ADORA2A* base edit (A_2A_R-KO) (**Supplementary Fig. 1**). A_2A_R-KO and unedited CAR-T cells were treated with the adenosine analogue 2-chloroadenosine (cADO) and evaluated for the level of phosphorylated cAMP Response Element-Binding Protein (pCREB), a downstream mediator of the adenosine-A_2A_R signaling pathway [41]. Notably, treatment with cADO increased the level of pCREB in unedited CAR-T cells, whereas the level of pCREB in A_2A_R-KO CAR-T cells did not deviate from the untreated control (**Fig. 1d,e**) indicating A_2A_R-KO attenuates proximal Ado-mediated signaling.

**Fig. 1:**
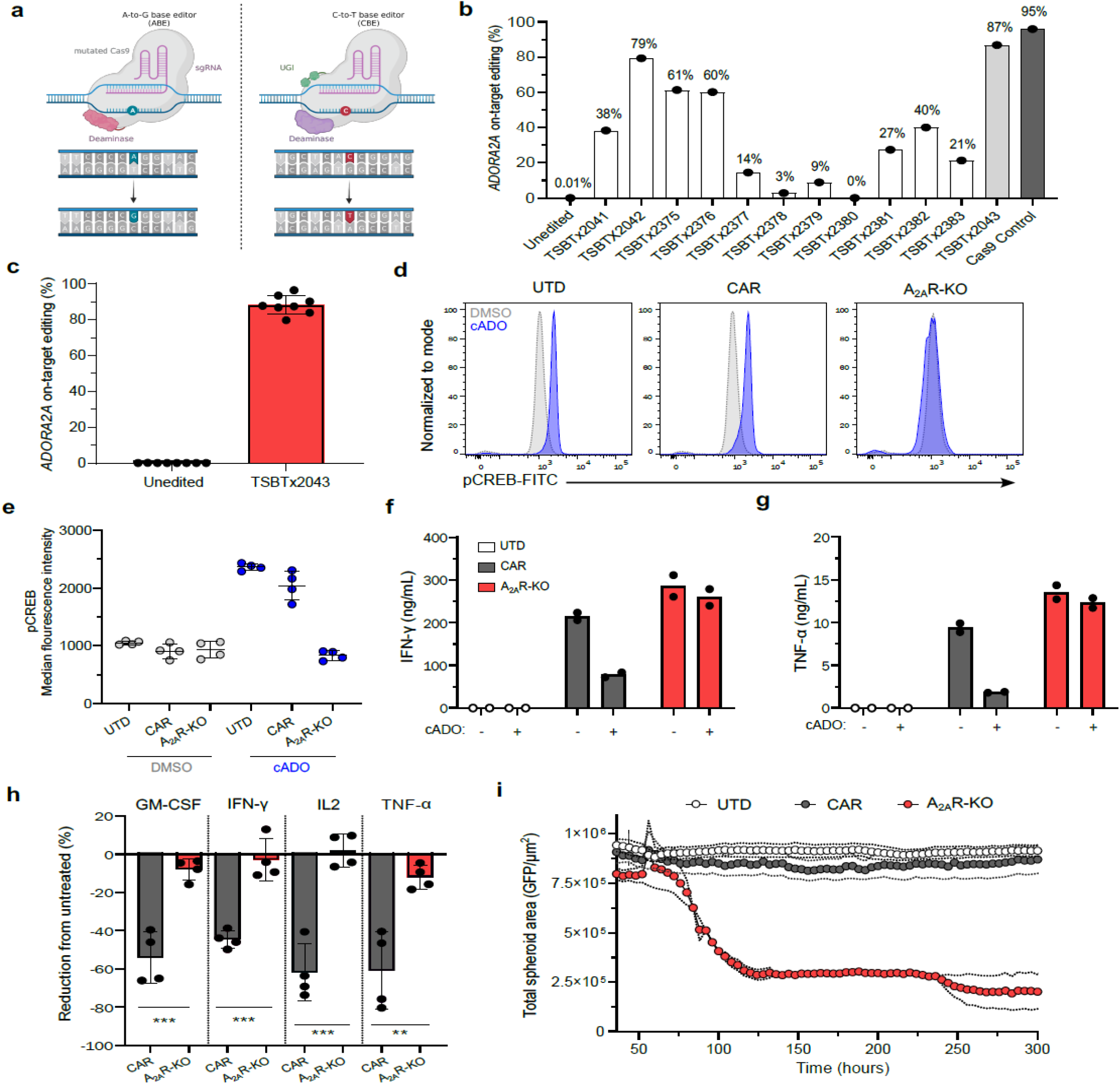
Base editing generates adenosine-resistant CAR-T cells. **a,** Schematic of adenine (ABE) and cytosine (CBE) base editor/DNA complexes, showing precise A-to-G (left) and C-to-T (right) base pair conversion. Mutated Cas9 protein (gray), single guide RNA (sgRNA, magenta), deaminase (left, pink and right, purple), uracil DNA glycosylase (UGI, green). **b,** Frequency of *ADORA2A* on-target genomic editing efficiency determined by next-generation sequencing (NGS) of each sgRNA-base editor pairing (ABE, light gray; CBE, white) in activated primary human T cells. **c,** Cumulative frequency of *ADORA2A* on-target genomic editing efficiency of top candidate sgRNA (TSBTx2043) complexed with ABE. **d,e,** Flow cytometry plots (**d**) and median fluorescence intensity (**e**) of phosphorylated CREB (pCREB) in untransduced (UTD) T cells, unedited CAR-T cells (CAR) and A_2A_R-edited CAR-T cells (A_2A_R-KO) treated with 2-chloroadenosine (cADO, blue) compared to an untreated control (DMSO, gray). **f,g,** IFN-γ (**f**) and TNF-α (**g**) production by UTD T cells (white), unedited (gray) and A_2A_R-KO (red) CAR-T cells measured by ELISA 48 hours post-stimulation with H226 tumor cells in the presence (+) or absence (-) of cADO. **h,** Magnitude of GM-CSF, IFN-γ, IL2, TNF-α secretion by unedited (gray) and A_2A_R-KO (red) CAR-T cells in the presence of cADO normalized to respective untreated CAR-T cells. **i,** Incucyte live imaging tumor spheroid cytotoxicity of UTD T cells (white), unedited (gray) and A_2A_R-KO (red) CAR-T cells. Cytotoxicity was measured as a decrease in cumulative tumor GFP^+^ area over time. For all data, bars and error bars reflect mean ± SD of individual biological replicates, except **f** and **g** where symbols represent technical replicates. **h**, Mann-Whitney t-test performed to calculate statistical significance, \*\*\**P*<0.001, \*\**P*<0.01.

We then investigated whether A_2A_R-KO alleviates Ado-induced immunosuppression by culturing CAR-T cells with EGFR^+^ lung cancer cell lines H226 or A549. After *in vitro* stimulation in the presence of cADO, A_2A_R-KO CAR-T cells maintained high levels of pro-inflammatory cytokine secretion including IFN-γ, TNF-α, IL2 and GM-CSF. In contrast, cADO potently suppressed cytokine production of unedited CAR-T cells relative to untreated CAR-T cells (**Fig. 1f-h; Supplementary Fig. 2**). Moreover, A_2A_R-KO CAR-T cells fully eradicated H226 tumor spheroids in the presence of cADO, whereas unedited CAR-T cells and untransduced (UTD) control T cells failed to exert tumor control (**Fig. 1i; Supplementary Fig. 3**). Together, these data demonstrate that base editor-mediated disruption of *ADORA2A* in CAR-T cells confers a functional resistance to Ado-mediated immunosuppression *in vitro*.

### A_2A_R-KO CAR-T cells overcome an immunosuppressive TME to control tumor progression *in vivo*

Ado accumulation in the TME is driven by hypoxia-responsive gene expression [42–44] thus, to interrogate the function of A_2A_R-KO CAR-T cells *in vivo*, we established a preclinical H226 xenograft mouse model that recapitulated a hypoxic TME. Immunofluorescent staining confirmed widespread hypoxia and expression of the Ado-generating ectoenzyme CD73 in H226 tumors resected from immunocompromised NCG mice (**Fig. 2a**). We then identified varying levels of hypoxia within the xenograft tumors based on the fluorescence intensity of hypoxia-sensitive pimonidazole (Hypoxyprobe) staining (**Fig. 2b**) and performed whole transcriptomic analysis using the Nanostring GeoMx digital spatial transcriptomics platform. Gene expression profiles of these regions were compared to the MGI Hypoxia Pathway Gene Ontology database, and notably, we observed induction of hypoxia -regulated genes, including *ALDOA, CA9, ENO1, LDHA, NDRG1, TGFB1* and *VEGF* proportional to the intensity of Hypoxyprobe staining (**Fig. 2c**). These data support the utility of this *in vivo* system to recapitulate immunosuppressive features of a hypoxic and Ado-rich TME.

**Fig. 2:**
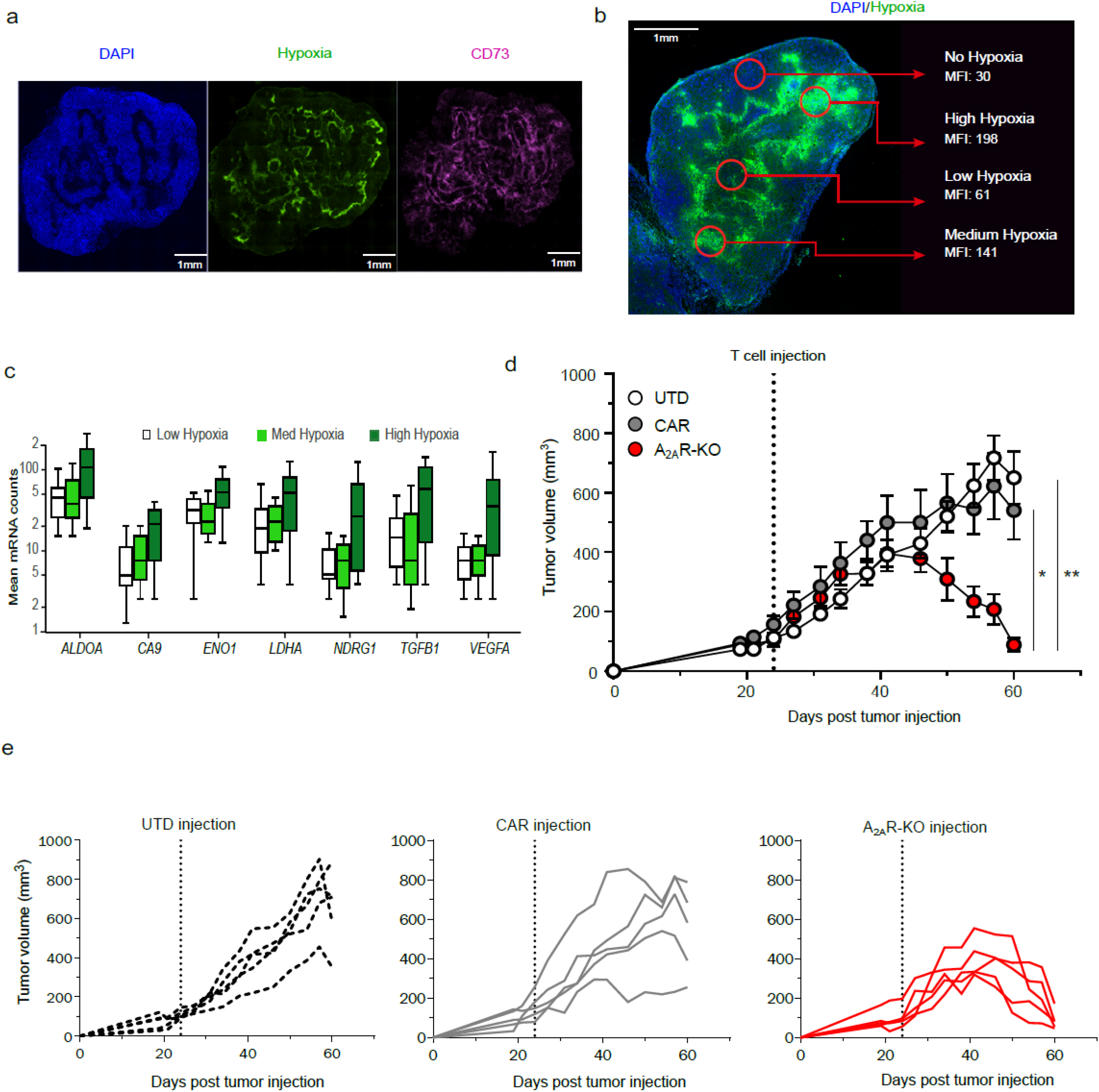
A_2A_R-KO CAR-T cells overcome hypoxia-adenosinergic suppression leading to improved tumor rejection *in vivo*. **a,** Representative immunofluorescent staining of H226 tumors resected from NCG mice 46 days post-implantation. Nucleated cells (DAPI, blue), hypoxia (Hypoxyprobe, green) and CD73 (pink). **b**, Quantification of hypoxia in tumor regions (red circles) within resected H226 tumors from NCG mice determined by mean fluorescence intensity (MFI) of Hypoxyprobe. **c,** Spatial transcriptomics gene expression analysis from low (white), medium (light green), and high (dark green) hypoxic regions in (**b**). **d,e,** 2x10^6^ UTD T cells (white, *n* = 5), unedited CAR-T cells (gray, *n* = 5) or A_2A_R-KO CAR-T cells (red, *n* = 5) were injected intravenously into H226 tumor bearing NCG mice once tumors achieved an average volume of 150mm^3^. Longitudinal group average (**d**) or individual mouse (**e**) tumor volumes measured using calipers. For all data, symbols and error bars reflect group mean ± S.E.M., except **c** where the bar represents median and box extends from the 25^th^ to the 75^th^ percentile for each group, and e where lines represent individual mice. **d**, Mann-Whitney t-test performed to calculate statistical significance, \*\**P*<0.01, \**P*<0.05.

To investigate whether elimination of A_2A_R enhances CAR-T cell activity *in vivo*, we treated H226 tumor bearing mice with three dose levels of A_2A_R-KO CAR-T cells, unedited CAR-T cells or UTD T cells. At medium and high dose levels, both A_2A_R-KO and unedited CAR-T cells controlled tumor outgrowth, but A_2A_R-KO CAR-T cell-treated mice exhibited lower peak tumor volume and accelerated kinetics of tumor regression (**Supplementary Fig. 4**). Notably, at the lowest evaluated dose, only A_2A_R-KO CAR-T cells fully eradicated tumors (**Fig. 2d,e**) and reduced cumulative tumor burden (**Supplementary Fig. 4**), while unedited CAR-T cells failed to control tumor. These findings demonstrate that A_2A_R-KO CAR-T cells overcome an immunosuppressive TME *in vivo* to suppress tumor progression.

### Alternative immunological mechanisms impair A_2A_R-KO CAR-T cell function

Hypoxic TMEs often comprise multiple immunosuppressive pathways including biochemical, metabolic, and immunological barriers that mitigate anti-tumor T cell functions [45,46]. Elimination of biochemical barriers alone, such as Ado, may not induce complete and durable tumor remissions [47](NCT02740985). Therefore, using digital spatial transcriptomics, we evaluated hypoxic regions in H226 xenografts and observed additional mechanisms of immunosuppression. Of note, regions of increasing hypoxia were associated with elevated levels of transforming growth factor beta 1 (TGF-β1) mRNA (**Fig. 2c**), which was then confirmed by detecting *in vitro* secretion of TGF-β by H226 tumor cells (**Supplementary Fig. 5**). In addition, flow cytometric analysis of cell surface proteins involved in T cell inhibitory pathways revealed that H226 tumor cells highly express PD-L1 (**Supplementary Fig. 5**). Indeed, both PD-L1 and TGF-β protein expression were confirmed by immunofluorescent staining of resected H226 tumors from NCG mice (**Fig. 3a**). We then tested whether A_2A_R-KO CAR-T cells were susceptible to PD-L1- or TGF-β-mediated inhibition. Upon *in vitro* stimulation in the presence of PD-L1, A_2A_R-KO CAR-T cells secreted lower amounts of IL-2 inversely proportional to the amount of PD-L1 inhibition (**Fig. 3b**). Similarly, the addition of exogenous TGF-β suppressed A_2A_R-KO CAR-T cell secretion of multiple pro-inflammatory cytokines including IL-2, IFN-γ, TNF-α and GM-CSF (**Fig. 3c, Supplementary Fig. 6**). These data indicate that A_2A_R-KO CAR-T cells remain sensitive to immunological negative regulators, and further suggest that PD-L1 and TGF-β may synergize with Ado in the TME to suppress anti-tumor T cell responses *in vivo*.

**Fig. 3:**
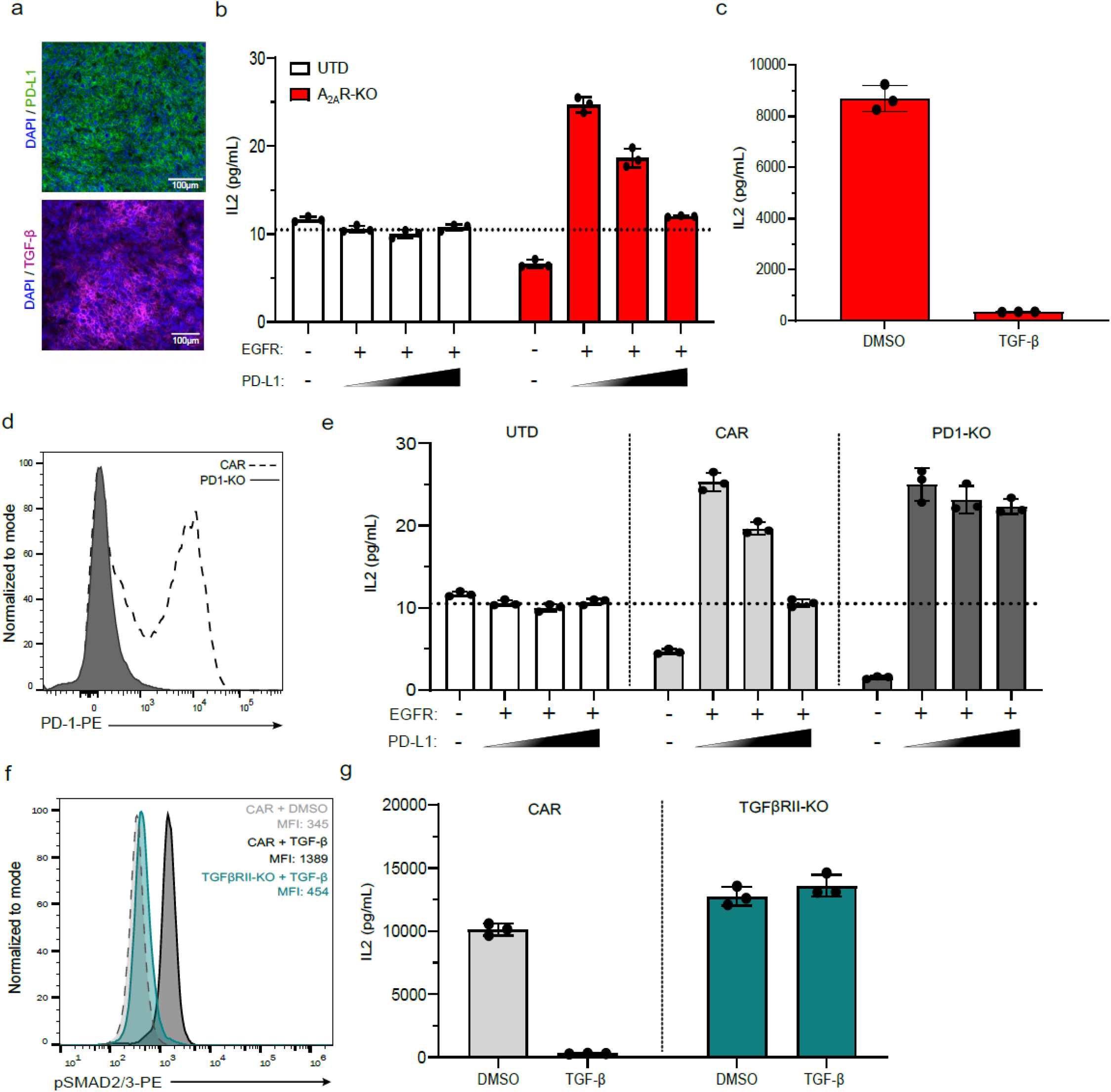
A_2A_R-KO CAR-T cells are susceptible to immunological negative regulators *in vitro*. **a,** Representative immunofluorescent staining of H226 tumors resected from NCG mice 46 days post-tumor implantation. Nucleated cells (DAPI, blue), PD-L1 (green), TGF-β (purple). **b,** IL2 production by UTD T cells (white) and A_2A_R-KO CAR-T cells (red) measured by ELISA 48 hours post-stimulation with recombinant human EGFR-conjugated Dynabeads and increasing amounts of PD-L-coated Dynabeads. **c**, IL2 production by A_2A_R-KO CAR-T cells 48 hours post-stimulation with H226 tumors cells in the presence of exogenous TGF-β or DMSO. **d,** Flow cytometry plot indicating PD-1 surface expression on unedited (dotted line) and PD1-KO (solid line) CAR-T cells 24 hours post-stimulation with PMA/Ionomycin. **e,** IL2 production by UTD T cells (white), unedited (light gray) and PD1-KO (dark gray) CAR-T cells measured 48 hours post-stimulation with recombinant human EGFR and PD-L1 conjugated beads. **f,** Flow cytometry plot of phosphorylated SMAD2/3 (pSMAD) in unedited (black) or TGFβRII-KO (green) CAR-T cells treated with exogenous TGF-β and unedited CAR-T cells treated with DMSO (dashed line). **g,** IL2 production by unedited (light gray) and TGFβRII-KO (dark gray) CAR-T cells 48 hours post-stimulation with H226 tumors in the presence of exogenous TGF-β. All functional assays were performed in duplicate, symbols represent technical replicates, and bars and error bars represent mean ± SD.

### Multiplex gene editing confers A_2A_R-KO CAR-T cells functional resistance to orthogonal immunosuppressive pathways

Since A_2A_R-KO CAR-T cells retained sensitivity to clinically relevant immunological barriers such as PD-L1 and TGF-β, we sought to simultaneously confer A_2A_R-KO CAR-T cells resistance to both inhibitory pathways. To do so, we utilized our previously described BE sgRNA targeting *PDCD1*, which encodes PD-1, the receptor responsible for PD-L1-mediated inhibition [36]. PD1-KO CAR-T cells did not upregulate PD-1 cell surface expression (**Fig. 3d**) and resisted PD-L1 mediated suppression of IL-2 secretion after *in vitro* antigen stimulation (**Fig. 3e; Supplementary 7**). Next, we evaluated sgRNAs spanning *TGFBR2* (**Supplementary Table 1**), the gene encoding TGF-β receptor 2 (TGFβRII) and screened each sgRNA-ABE complex in primary human T cells. None of the evaluated sgRNAs drastically attenuated downstream TGF-β-mediated signaling indicated by the phosphorylation of SMAD2/3 (**Supplementary Fig. 8**). Therefore, we designed CRISPR-Cas12b nuclease sgRNAs, and with its unique ATTN protospacer adjacent motif (PAM) sequence, were able to target alternative loci within *TGFBR2* compared to ABE (**Supplementary Table 1**). In doing so, we identified a Cas12b-sgRNA pairing that completely abrogated pSMAD2/3 relative to unedited control T cells (**Fig. 3f**), and after *in vitro* tumor stimulation, TGFβRII-KO CAR-T cells maintained secretion of effector cytokines despite the presence of exogenous TGF-β (**Fig. 3g; Supplementary Fig. 9**).

Next, we co-introduced three genetic edits, termed triple knock-out (TKO), into EGFR-specific CAR-T cells to simultaneously overcome Ado, PD-L1 and TGF-β tumor-associated inhibitory pathways (**Fig. 4a**). TKO CAR-T cells were generated by combining *ADORA2A-*, *PDCD1-,* and *TGFBR2*-specific sgRNAs into a single electroporation with mRNAs encoding ABE and Cas12b, which achieved genomic on-target editing efficiencies of 93% ± 2.5%, 95% ± 1.7%, and 92% ± 3.8%, respectively (**Fig. 4b**). To determine whether multiplex gene editing confers functional resistance to these biochemical and immunological inhibitory pathways, TKO CAR-T cells were tumor stimulated in an *in vitro* system that concomitantly recapitulates Ado, PD-L1 and TGF-β inhibition. TKO CAR-T cells resisted the suppressive effects of all three mechanisms as evidenced by their ability to maintain elevated levels of pro-inflammatory cytokine secretion (**Fig. 4c, Supplementary Fig. 10**). In stark contrast, the anti-tumor activity of unedited CAR-T cells, A_2A_R-KO CAR-T cells, and even CAR-T cells containing both PD1-KO and TGFβRII-KO (termed double knockout, DKO) were attenuated by these inhibitory pathways (**Fig. 4c; Supplementary Fig. 10**). Notably, using the same *in vitro* suppression system, TKO CAR-T cells fully eradicated H226 tumor spheroids, exhibiting superior anti-tumor activity compared to A_2A_R-KO CAR-T cells (**Fig. 4d**). These findings demonstrate that multiplex gene editing simultaneously confers TKO CAR-T cells functional resistance to multiple immunosuppressive pathways *in vitro*.

**Fig. 4:**
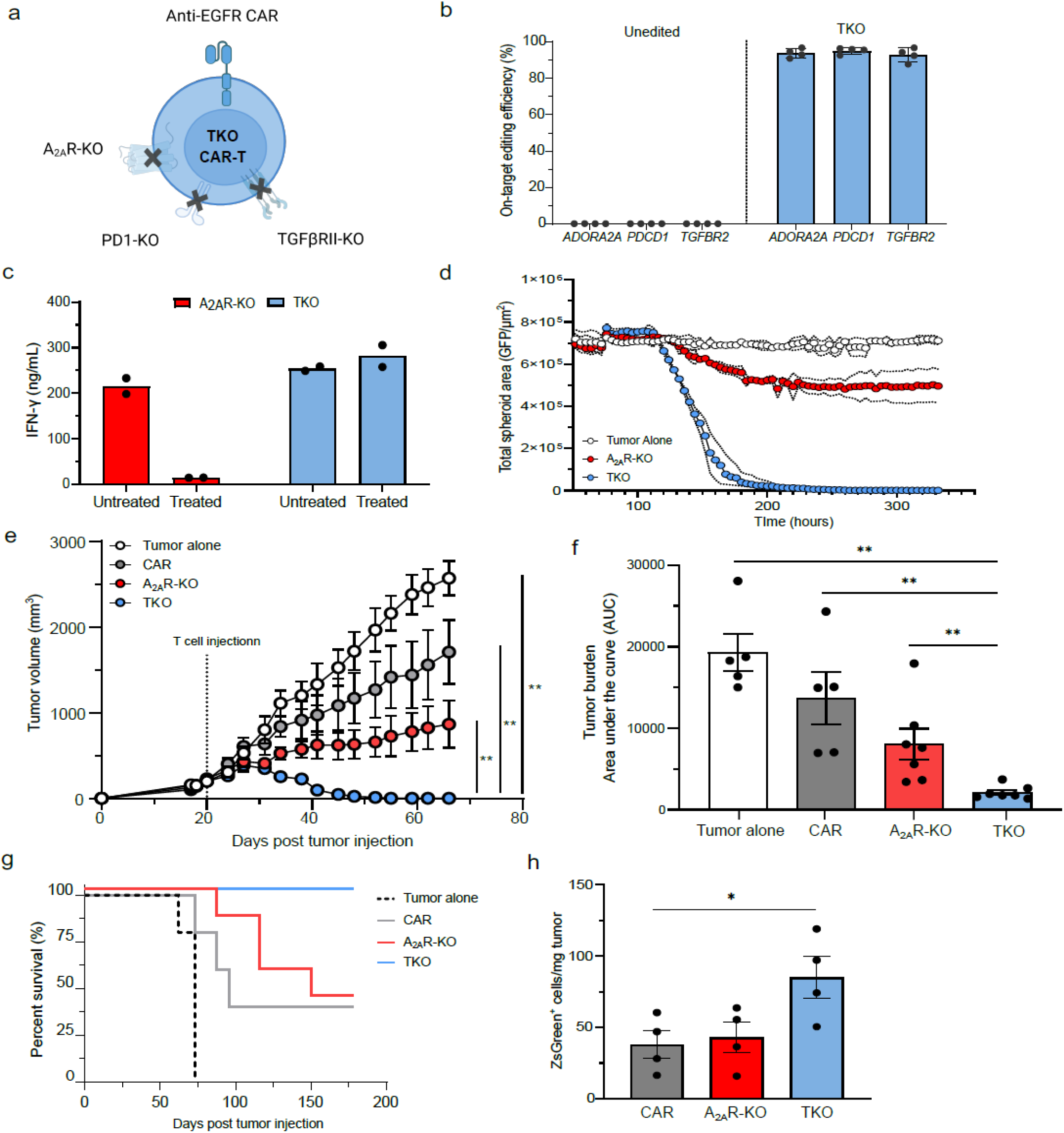
Comprehensive genome editing confers CAR-T cells resistance to multiple inhibitory pathways. **a,** Schematic of triple knock out (TKO) EGFR-specific CAR-T cells comprising three genomic edits targeting *ADORA2A* (A_2A_R-KO), *PDCD1* (PD-1-KO) and *TGFBR2* (TGFβRII-KO). **b,** Frequency of *ADORA2A*, *PDCD1*, and *TGFBR2* on-target genomic editing efficiencies in unedited and TKO CAR-T cells quantified by NGS. **c,** IFN-γ production by A_2A_R-KO (red) and TKO (blue) CAR-T cells measured by ELISA 48 hours post-stimulation with H226 tumors in the presence of DMSO (untreated) or triple suppression comprising exogenous cADO, PD-L1 and TGF-β proteins (treated). **d,** Incucyte-based live imaging of GFP^+^ H226 tumor spheroid cytotoxicity of tumor alone (white) and tumor cultured with A_2A_R-KO (red) or TKO (blue) CAR-T cells in the presence of triple suppression. Cytotoxicity was measured as a decrease in total tumor GFP^+^ area over time. **e-h,** NCG mice were implanted with H226 tumor and untreated (tumor alone, dotted line, *n* = 5) or were injected with 2x10^6^ ZsGreen^+^ unedited (gray, *n* = 5), A_2A_R-KO (red, *n* = 7) or TKO (blue, *n* = 7) CAR-T cells when tumors reached an average volume of 150mm^3^. Longitudinal tumor volume (**e**), cumulative tumor burden calculated as area under the curve (**f**), and survival curves (**g**) are shown. **h,** Summary data enumerating total ZsGreen ^+^ CAR-T cells per milligram of resected H226 tumors from NCG mice 7 days post-T cell infusion. For all data, bars and error bars reflect mean ± SD and symbols represent individual mice, except **c** where symbols are technical replicates, and **e,** where error bars represent S.E.M. **f**,**h**, Kruskal-Wallis test performed to calculate statistical significance, \*\**P*<0.01, \**P*<0.05.

To determine if TKO CAR-T cells overcome the convergence of these negative regulators *in vivo*, we compared their potency to A_2A_R-KO CAR-T cells and unedited CAR-T cells at subtherapeutic dose levels in H226 tumor bearing NCG mice. As expected, unedited CAR-T cells demonstrated limited capacity to control tumor outgrowth, while A_2A_R-KO CAR-T cells mitigated tumor progression but did not clear tumor. However, TKO CAR-T cells durably eradicated established tumors (**Fig. 4e; Supplementary Fig. 11**), decreased cumulative tumor burden (**Fig. 4f**), and extended the lifespan of treated mice (**Fig. 4g**). In a parallel mechanistic study, tumors were excised from CAR-T cell treated mice 1-week post-infusion to quantify T cell infiltration in the TME. TKO CAR-T cells were more abundant in the TME compared to A_2A_R-KO and unedited CAR-T cells (**Fig. 4h; Supplementary Fig. 12**), likely owing to their ability to penetrate and locally proliferate within the immunosuppressive TME. Together, these data highlight the power of multiplex gene editing to generate a CAR-T cell product that is simultaneously resistant to multiple TME-associated immunosuppressive features *in vivo*.

### Stealth TKO CAR-T cells resist allorejection in immunocompetent mice and potently reject tumors

A limitation of tumor-engrafted immunocompromised mice is the paucity of immunologically relevant human cell types that contribute to forming the TME. Therefore, we wanted to investigate the functionality of TKO CAR-T cells in a small-animal model that reconstitutes a human immune system. To do so, we utilized humanized NCG (huNCG) mice that are generated via adoptive transfer of human cord blood-derived CD34^+^ hematopoietic stem cells into recipient mice. huNCG mice developed peripheral CD4^+^ and CD8^+^ T cells, B cells, myeloid cells, and a limited pool of mature NK cells (**Supplementary Fig. 13**). Importantly, CD4^+^FoxP3^+^ regulatory T cells and CD68^+^ tumor-associated macrophages were detected in resected tumor sections from H226 bearing huNCG mice (**Supplementary Fig. 14**) indicating that these mice recapitulate a tumor milieu resembling TMEs described in humans [48–51].

Additionally, successful engraftment of adoptively transferred allogeneic CAR-T cells into a genetically dissimilar lymphoreplete host relies on incorporating a cloaking strategy to evade the host immune system. Given that huNCG mice primarily reconstitute a human T cell compartment, we used ABE to simultaneously introduce two ‘Stealth’ gene edits into allogeneic CAR-T cells targeting *B2M* (beta-2-microglobulin) and *CIITA* (class-II transcriptional activator) to disrupt HLA class-I and HLA class-II surface expression thereby preventing recipient CD8^+^ and CD4^+^ T cell-mediated rejection, respectively. Additionally, these HLA-deficient CAR-T cells were edited for *CD3E* to ablate surface expression of the endogenous T cell receptor complex preventing graft-versus-host disease (**Fig. 5a**). To determine whether genetic disruption of *B2M* and *CIITA* protect allogeneic donor-derived CAR-T cells from immune-mediated rejection, we co-transferred allogeneic HLA-deficient (Stealth) and unedited HLA^+^ CAR-T cells into recipient huNCG mice, and the persistence of these cell products was monitored by longitudinally sampling peripheral blood. Unedited CAR-T cells were readily eliminated from peripheral blood within 2 weeks post-infusion, whereas Stealth CAR-T cells resisted allorejection and persisted for the duration of the study (**Fig. 5b,c**).

**Fig. 5:**
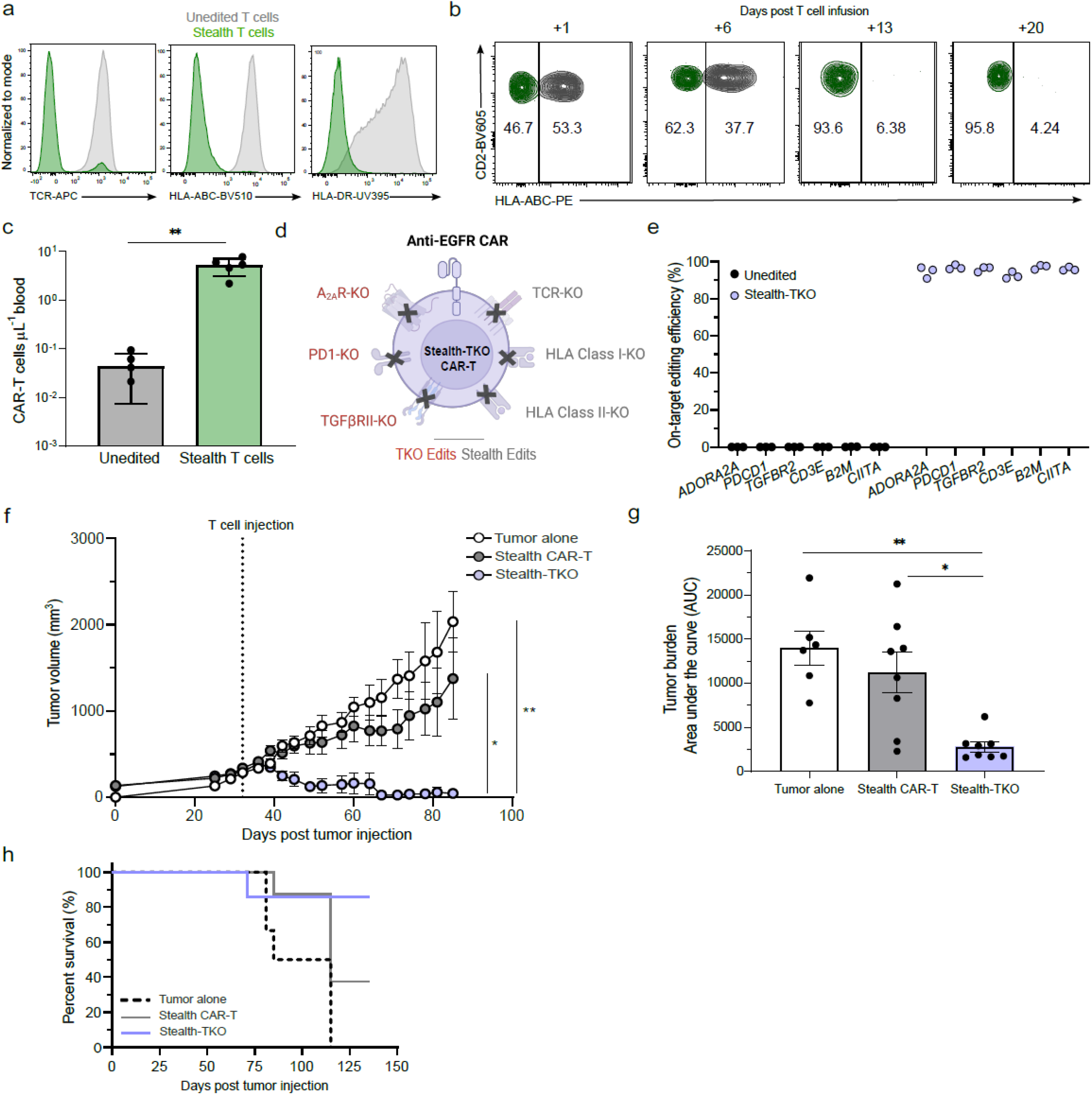
Stealth-TKO CAR-T cells resist allorejection and clear tumor in humanized mice. **a,** Flow cytometry histograms indicating surface expression of endogenous TCR, HLA-class I (HLA-ABC) and HLA-class II (HLA-DR) on unedited (gray) and Stealth (green) T cells. **b, c,** 5x10^6^ unedited (gray, *n* = 4) and Stealth (green, *n* = 5) CAR-T cells were co-infused into humanized NCG mice. Flow cytometry plots indicate longitudinal frequency of peripheral allogeneic T cells from within the same mouse at 1-, 6-, 13- and 20-days post-infusion (**b**) and cumulative blood concentration of peripheral T cells 20 days post-infusion (**c**). **d,** Schematic of Sealth-TKO EGFR-specific CAR-T cells comprising six genomic edits targeting *ADORA2A* (A_2A_R-KO), *PDCD1* (PD-1-KO), *TGFBRII* (TGFBR2-KO), *CD3E* (TCR-KO), *B2M* (HLA-Class I-KO), and *CIITA* (HLA-Class II-KO). **e,** Frequency of on-target genomic editing efficiencies at six genomic loci in Unedited (black) and Stealth-TKO (purple) CAR-T cells quantified by NGS. **f-h,** H226 tumor bearing hNCG mice were untreated (white, *n* = 6) or were infused with 4x10^6^ Stealth CAR-T cells (gray, *n* = 8) or Stealth-TKO CAR-T cells (purple, *n* = 8) when tumor volume reached an average volume of 150mm^3^. Longitudinal tumor volume (**f**), cumulative tumor burden (**g**) and survival curves (**h**) are shown. For all data, symbols represent individual mice or T cell donors, and error bars reflect mean ± SD, except **f** where error bars represent S.E.M. **c**, Mann-Whitney and **g**, Kruskal-Wallis test performed to calculate statistical significance, \*\**P*<0.01, \**P*<0.05.

After determining that Stealth CAR-T cells successfully engraft into huNCG mice, we engineered TKO CAR-T cells with *CD3E*, *B2M* and *CIITA* base edits (**Fig. 5d**). The resulting 6-plex gene edited cell product, termed Stealth-TKO CAR-T cells, maintained a high degree of on-target genomic editing efficiency at all loci (**Fig. 5e**). Upon infusion into H226 tumor bearing huNCG mice, Stealth -TKO CAR-T cells potently eliminated tumor at multiple dose levels (**Fig. 5f; Supplementary Fig. 15**). In contrast, the tumor growth kinetics in mice treated with CAR-T cells containing only Stealth edits mirrored untreated control mice (**Fig. 5f**). Moreover, the superior anti-tumor responses mediated by Stealth-TKO CAR-T cells significantly reduced cumulative tumor burden and extended the overall lifespan of tumor bearing mice (**Fig. 5g,h**). The ability of Stealth-TKO CAR-T cells to evade allorejection and mediate potent anti-tumor activity despite a complex TME within a lymphoreplete host underscores the utility of combinatorial gene editing to derive CAR-T cell therapies with great translational promise.

## Discussion

Overcoming multiple immunosuppressive barriers acting on CAR-T cells within the TME likely requires a multifaceted genome engineering strategy. To this end, we employed a novel combination of our Base Editing platform [31] with a CRISPR nuclease [37] to manufacture allogeneic ‘off-the-shelf’ CAR-T cells resistant to biochemical and immunological negative regulators. These data emphasize the importance of understanding CAR-T cell-TME interactions, as well as the therapeutic advantages of simultaneously blocking multiple immunosuppressive pathways.

Solid tumors promote hypoxic and adenosine-rich microenvironments that can be ameliorated by pharmacologic inhibitors [52–55]. While initial Phase II (NCT05024097) and Phase III (NCT05221840) clinical results are promising, antagonists of the adenosinergic pathway may be ineffective due to the presence of alternative suppressive mechanisms and the paucity of anti-tumor T cells in the TME. To ensure sufficient numbers of tumor-reactive T cells, we developed solid tumor-specific CAR-T cells that are resistant to adenosine-mediated immunosuppression. We demonstrated the first use of base editing in CAR-T cells to genetically ablate *ADORA2A*, the gene encoding A_2A_R, which is the major receptor that binds extracellular adenosine initiating inhibitory programming. Ablation of A_2A_R in CAR-T cells attenuated downstream phosphorylation of CREB, increased production of pro-inflammatory cytokines and improved tumor elimination, in accordance with observations by Beavis et al. using CRISPR-Cas9 mediated elimination [30]. However, A_2A_R-KO CAR-T cells retained sensitivity to immunological negative regulators such as PD-L1 and TGF-β. Likewise, CAR-T cells deficient for both PD-1 and TGF-βRII were highly suppressed by adenosinergic signaling. These data support the notion that i) non-redundant inhibitory mechanisms within the TME act in concert to inhibit anti-tumor responses, and ii) engineering CAR-T cells to overcome a single pathway is likely insufficient to engender durable remissions.

Here, we demonstrated that simultaneous disruption of biochemical (adenosine) and immunological (PD-L1 and TGF-β) inhibitory pathways is critical to maximize the therapeutic efficacy of solid tumor-specific CAR-T cells. Recent research has identified additional negative regulators within the TME [56–58] and is likely to reveal more target genes for interrogation, ultimately requiring a complex genome engineering strategy to generate higher-order multiplex gene-edited CAR-T cells. To meet the challenge of complex genome engineering, we combined an adenine base editor (ABE) and Cas12b, a nuclease with a distinct ATTN protospacer adjacent motif (PAM) sequence dissimilar to the NGG PAM used by ABE [37]. By utilizing multiple PAMs, this unique combination of editors complementarily acts to expand the range of accessible target sites up to 6 genes reported here and provides future avenues for nuclease-driven gene knock-in. We believe this strategy provides proof-of-concept rationale for the design of CAR-T cell products with a synergistic combination of gene edits that confer potent anti-tumor responses.

Moreover, we have demonstrated the power of multiplex gene editing in successfully incorporating additional edits to manufacture an allogeneic ‘off-the-shelf’ CAR-T cell product. CAR-T cells derived from an allogeneic donor hold promise in solving challenges associated with autologous, patient-derived products including manufacturing failure and the urgency to reduce patient wait-time to treatment [59–62]. Thus, a critical aspect of this study applied ‘Stealth’ gene edits to TKO CAR-T cells to prevent GvHD (CD3E-KO) and evade T cell-mediated rejection caused by HLA haplotype mismatch between patient and allogeneic CAR-T cell donor (B2M-KO and CIITA-KO). Disrupting these target genes permitted stable engraftment of allogeneic CAR-T cells into lymphoreplete humanized mice resulting in superior solid tumor clearance. However, it is important to note that stem cell -engrafted humanized mice often exhibit incomplete immune reconstitution, e.g., mature NK cells [63], suggesting that additional modifications such as genetic ablation of NK cell activating receptors [64] or overexpression of the invariant HLA-E inhibitory receptor [65] may be necessary to truly render allogeneic CAR-T cells hypoimmunogenic.

This study provides rationale and precedent for the use of combinatorial gene editing to manufacture an allogeneic ’off-the-shelf’ CAR-T cell product that maximizes therapeutic efficacy by overcoming orthogonal immunosuppressive barriers in the solid TME. While it must be considered that an inflection point may be reached whereby increasing the number of edits decreases overall CAR-T cell viability or functionality, our data indicate that this was not the case with the 6-plex edited Stealth-TKO CAR-T cells. By employing a unique Base Editor/CRISPR nuclease combination, we focused on eliminating biochemical and immunological barriers that are common to many solid TMEs in an effort to enhance the therapeutic capacity of CAR-T cells. Thus, the engineering approach described here may unlock the full potential of solid tumor targeting CAR-T cells.

## Materials and Methods

### Cell lines

NCI-H226, A549, H460, MDA-MB-231, SKOV3, MCF7, A375 cells were obtained from the American Type Culture Collection (ATCC). NCI-H226 and H460 cells were grown in RPMI-1640 medium (Gibco) supplemented with 10% fetal bovine serum (Gibco) and A549, MDA-MB-231, SKOV3, MCF7 and A375 cells were grown in DMEM medium (Gibco) supplemented with 10% fetal bovine serum at 37°C in a 5% CO_2_ incubator.

### Mouse strains and study approval

Female 6–8-week-old NOD-*Prkdc^em26Cd52^Il2rg^em26Cd22^*/NjuCrl (NCG, strain code: 572) coisogenic immunodeficient mice and human umbilical cord blood-derived CD34^+^ humanized coisogenic NOD-*Prkdc^em26Cd52^Il2rg^em26Cd22^*/NjuCrl female mice (huNCG, strain code: 695) mice were obtained from Charles River Laboratories. HuNCG received 12-15 weeks post-humanization. Animals were housed in a specific pathogen–free environment under controlled conditions and received food and water ad libitum according to the National Institutes of Health (NIH) guidelines. All animal experiments were conducted in accordance with the guidelines and approval of the Institutional Animal Care and Use Committee at Northeastern University and the Charles River Accelerator and Development Lab.

### Flow cytometry

Surface expression of anti-EGFR CARs was detected by staining with an anti-Cetuximab-AF647 idiotype (Clone: 2259C, R&D Systems). T cell editing efficiency, T cell phenotyping, T cell signaling, tumor cell phenotyping and mouse blood phenotyping were evaluated with the following anti-human antibodies: CD155 (SKII.4, Biolegend), CD19 (HIB19, Biolegend), CD2 (RPA-2.10, Biolegend), CD3 (UCHT1, Biolegend), CD33 (P67.6, Biolegend), CD38 (HIT2, Biolegend), CD39 (A1, Biolegend), CD4 (OKT4, Biolegend), CD45 (2D1, Biolegend), CD47 (CC2C6, Biolegend), CD56 (NCAM, Biolegend), CD68 (FA11, Biolegend), CD73 (AD2, Biolegend), CD8 (SK1, Biolegend), CD80 (B7 -1, BD Biosciences), CD86 (IT2.2, Biolegend), EGFR (AY13, Biolegend), CD95L (NOK-1, Biolegend), HLA-ABC (W6/32, Biolegend), PD1 (A17188B, Biolegend), PDL1 (29E.2A.3, Biolegend), TCR (IP26, Biolegend). CD73 (AD2, Cell Signlaing), PDL1 (D8T4X, Cell Signaling), Phospho -CREB (87G3, Cell Signaling), HLA-DR (L203.rMAb, BD Biosciences), Phospho-SMAD2/3 (027-670, BD Biosciences), TGFβ1 (Polyclonal, Bioss). Antibodies for Mass Cytometry all supplied from Fluidigm: Alpha-SMA (1A4), Collagen-1 (polyclonal), E-cadherin (2.4e+11), Histone H3 (D1H2), Vimentin (D21H3), Granzyme B (EPR20129-217), Ki-67 (B56), PD-1 (EPR4877(2)), PD-L1 (Sp142), CD20 (H1), CD3 (polyclonal), CD4 (EPR6855), CD45RO (UCHL1), CD68 (KP1), CD8α (CD8/144B), FoxP3 (PCH101), Pan-keratin (C11). For detection of surface proteins, cells were stained for 20 minutes at 4°C in the dark. For intracellular staining of phosphorylated proteins, cells were stimulated with agonist and then fixed with pre-warmed Cytofix (BD Biosciences) for 15 minutes at 37°C, then permeabilized with Perm III buffer (BD Biosciences) for 30 minutes at -20°C. Fixed and permeabilized cells were then stained with antibodies for 1 hour at room temperature in the dark. All samples were washed with PBS prior to analysis on LSR Fortessa (BD Biosciences) flow cytometer and data analyzed using FlowJo v.10.9 software.

### Generation of CAR-T cells

CD4^+^ and CD8^+^ cells were positively selected from a de-identified healthy human donor apheresis (Charles River Laboratories) using anti-CD4 and anti-CD8 microbeads (Miltenyi) following the manufacturer’s protocol. Post isolation, cells were washed and cryopreserved in a 1:1 mixture of CS10 (BioLife Solutions) and Plasma-Lyte A (Hanna Pharmaceutical) supplemented with 2% HSA (Access Biologicals). Cryopreserved cells were thawed and activated with anti-CD3/anti-CD28 TransAct (Miltenyi) 1:100 v/v in T cell expansion medium comprising OpTmizer media (Thermo Fisher) containing 2% GlutaMAX (Thermo Fisher), 2.5% Immune cell serum replacement (Thermo Fisher), 100 IU/mL rhIL-2, 2ng/mL rhIL-7 and 0.4ng/mL rhIL-15 (R&D Systems). T cells were transduced 24 hours post-activation with lentivirus encoding an anti-EGFR CAR with 41bb/CD3ζ, signaling domains, or co-transduced with an additional lentivirus encoding ZsGreen protein where applicable (Flash Therapeutics). 48 hours post-activation, T cells were prepared for gene editing. Briefly, T cells were washed with PBS and resuspended at 100x10^6^ cells/mL in Lonza P3 Primary Cell 4D Nucleofector Kit buffer (Lonza). 1μg/1x10^6^ cells of sgRNA (Agilent) (**Supplementary Table 1**) and corresponding 1μg/1x10^6^ of editor mRNA encoding ABE, CBE or Cas12b were added to P3 buffer containing T cells. T cells were electroporated using the Lonza 4D Nucleofector X Unit. Post-electroporation, cells were transferred to GRex culture flasks (Wilson Wolf) and expanded at 37°C in a 5% CO_2_ incubator. Verification of editing was confirmed via flow cytometry and next generation sequencing 10 days post T cell activation.

### Next generation Sequencing

Genomic DNA was extracted from cell pellets with QuickExtract DNA Extraction Solution (Lucigen) via manufacturer’s protocol. Subsequently, 2μl of genomic DNA was added to a 25μl PCR reaction containing Phusion U Green Multiplex PCR Master Mix (Thermo Fisher) and 0.5μM of each forward and reverse primer for each target site. Following initial PCR amplification, PCR products were barcoded using Illumina barcoding primer pairs. DNA was sequenced on an Illumina MiSeq instrument according to the manufacturer’s protocol. NGS data was analyzed by first performing Illumina demultiplexing, then read trimming and filtering, followed by alignment of all reads to the reference amplicon sequence prior to quantification generation of alignment statistics and editing rates.

### A_2A_R signaling assay

CAR-T cells were placed at 37°C in a 1% O_2_ incubator for 48 hours. Following hypoxic pretreatment, cells were washed in PBS and counted prior to assay. 1x10^6^ cells were plated per well of 96-well round bottom plate. Cells were treated with 30µM 2-chloroadenosine (cADO, Sigma) or equivalent volume of DMSO for 1 hour at 37°C. Subsequently, cells were stained for phosphorylated CREB and analyzed as described above.

### TGFβR signaling assay

CAR-T cells were rested in media without serum replacement overnight at 37°C in a 5% CO_2_ incubator. Following serum starvation, cells were washed in PBS and counted prior to assay. 1x10^6^ cells were plated per well of 96-well round bottom plate and treated with 10ng/mL of TGF-β1 (Peprotech) or equivalent volume of DMSO for 20 minutes at 37°C. Cells were then stained for phosphorylated SMAD2/3 and analyzed as described above.

### Cytokine Release Assay

1x10^5^ H226 or A549 tumor cells and 5x10^4^ CAR^+^ T cells were plated per well of 96-well flat bottom plate. DMSO, 10µM cADO or 10ng/mL TGF-β1 were added, and cells were incubated at 37°C, 5% CO_2_ for 48 hours. At assay endpoint, plates were centrifuged at 500xg for 5 minutes, and supernatants taken for ELISA analysis of GM-CSF, IL2, IFN-γ, and TNF-α on the Ella platform (Protein Simple) following the manufacturer’s protocol.

### PD-1 induction assay

1x10^6^ T cells were plated per well of 96-well round bottom plate. Cells were treated with 1:1000 diluted Cell Stimulation cocktail (Biolegend) overnight at 37°C, 5% CO_2_. Next, cells were washed with PBS, and stained with anti-PD1 antibody as described above. Data acquired on LSR Fortessa (BD Biosciences) and analyzed via FlowJo software.

### PD-L1 Bead Suppression Assay

Saturating amounts of recombinant human (rh) EGFR-Biotin and rhPD-L1-Biotin (ACRO Biosystems) were coated onto separate Streptavidin-coated DynaBeads (Thermo Fisher) using manufacturer’s protocol. Protein-labeled beads were washed twice in PBS and resuspended in T cell expansion media at 6.5x10^8^ beads/mL. 1x10^6^ CAR-T cells were plated per well of 96-well flat bottom plate. 2µL of EGFR-coated beads and either 2µL, 5µL or 10µL of PD-L1-coated beads were added to each well and incubated at 37°C, 5% CO_2_ for 48 hours. At assay endpoint, plates were centrifuged at 500xg for 5 minutes, and supernatants were taken for ELISA analysis of IL2 on Ella platform via manufacturer’s protocol.

### Cytotoxicity Assay

H226 and A549 lung carcinoma cell lines were stably transduced with lentivirus encoding GFP. 1.5x10 ^4^ GFP^+^ tumor cells were plated in ultra-low attachment 96-well plates (Corning) and placed at 37°C, 5% CO_2_ for 72 hours to allow for spheroid formation. After 72 hours, 7x10^3^ CAR-T cells were plated in tumor wells with or without 10µM cADO, 10ng/mL TGF-β1 or 10µL PD-L1 coated beads. Cytotoxicity was quantified as reduction of GFP^+^ area over time via imagining in Incucyte S3 Live-Cell Analysis System (Sartorius).

### *In vitro* tumor cell TGF-β production assay

1x10^6^ NCI-H226, A549, H460, MDA-MB-231, SKOV3, MCF7 or A375 tumor cells were plated in 24-well tissue culture plate in 1mL of media. After 24 hours, supernatants were taken for ELISA analysis of TGF-β1 on Ella platform via manufacturer’s protocol.

### CAR-T cell efficacy *in vivo*

5x10^6^ H226 cells were resuspended in 200μL PBS and injected subcutaneously into the hind limb of NCG or huNCG mice. Tumor size was measured via calipers 2-3 times per week. Once tumors achieved an average volume of 150mm^3^, mice were injected with either 2x10^6^, 4x10^6^ or 8x10^6^ CAR-T cells. Control mice were either untreated (tumor only) or infused with 2x10^6^, 4x10^6^ or 8x10^6^ untransduced (UTD) T cells. T cells were injected via tail vein in 200µL HBSS. Weekly submandibular bleeds were used to track human cells within the blood over the course of the experiment. Prior to sacrificing the animals, mice were injected with 80mg/kg of Hypoxyprobe (pimonidazole HCl, 4 .3.11.3, Hypoxyprobe) solution via tail vein. 60 minutes post-Hypoxyprobe injection, the mice were sacrificed, and tumors resected. Tumors were washed in PBS and then flash frozen in OCT blocks and stored at -80°C.

### Allogeneic CAR-T cell *in vivo* persistence assay

CD2^+^CD3^+^HLA-ABC^+^HLA-DR^+^ (unedited) and CD2^+^CD3^-^HLA-ABC^-^HLA-DR^-^ (Stealth) T cells expressing a non-targeting CD4-based CAR were generated as previously described [66]. 5x10^6^ CAR-T cells of each population were co-injected intravenously into huNCG mice. Blood was collected via puncture of the submandibular vein 1-day post-infusion and weekly thereafter until 20 days post-infusion, into K2 EDTA coated microvette tubes (Sarstedt Inc) and stained as described above and analyzed on MACSQuant 16 (Miltenyi) flow cytometer.

### Tissue Sectioning and Immunohistochemistry

Tumors were excised from mice, washed in PBS, flash frozen into OCT blocks and stored at -80°C until further processing. Blocks were removed from freezer and placed in pre-chilled NX70 CryoStat (ThermoFisher). 5μm tissue sections were mounted onto polysine-coated slides and air-dried for 45-60 mins. Sections were fixed in a 1:1 mixture of acetone and methanol for 10 minutes and subsequently air-dried for 10 mins. Next, hydrophobic barrier PAP pen was applied to the edges of the sections and IHC buffer (PBS, 0.5% BSA, 0.1% Tween-20) added for 20 mins. Then, sections were blocked with Fc block in IHC buffer for 10 minutes followed by immunostaining of antibody cocktail in IHC buffer for 3 hours at room temperature. Slides were then washed three times with IHC buffer for 5 minutes, stained with DAPI, coverslips applied with fluoromount and imagined with an Olympus IX83 inverted microscope and analyzed using the Olympus CellSense software.

### Detection of tumor infiltrating CAR-T cells *ex vivo*

ZsGreen^+^ CAR-T cells were intravenously injected into H226 tumor-bearing mice. Tumors were then excised 7-days post T cell infusion, washed in PBS and transferred to a 50mL conical tube containing RPMI-1640 media and digestion enzymes from the human tumor dissociation it (Miltenyi). Tumors were then digested using Miltenyi GentleMACS tissue dissociator accoring to the manufacturer’s protocol. Following digestion, tumors were centrifuged at 300g for 5 minutes and supernatant aspirated. Cell pellets were resuspended in Percoll and centrifuged at 300g for 10 minutes. Supernatant was again aspirated prior to the addition of 1mL ACK lysis buffer (Thermo Fisher). Cells were incubated in buffer for 5 minutes at room temperature. Finally, 5mL of RPMI-1640 media containing serum was added, and cells centrifuged at 500g for 5 minutes. After supernatant was aspirated, cells were prepped and stained as described above and tumor infiltrating CAR-T cells were analyzed using LSR Fortessa.

### Spatial Transcriptomics

Tumors were excised and prepared as described for immunohistochemistry analysis. Tissues were mounted on slides, prepared and stained per Nanostring protocol. Briefly, tissue sections were fixed with 10% NBF overnight. Next, slides were washed in PBS, rehydrated in ethanol, treated with proteinase K and hybridized with RNA probes from the Nanostring Human Whole Transcriptome Atlas panel overnight at 37°C. Next, slides were washed and stained with nuclear dye Syto-83 or DAPI (Thermo Fisher), PD-L1 (Cell Signaling), TGFβ1 (Bioss), Hypoxyprobe, or CD73 (Cell Signaling) for 1 hour at room temperature. Slides then washed and imaged on GeoMX DSP (Nanostring). RNA was collected and run on the MiSeq (Illumina). Standard quality control checks assessing imaging, binding density, positive control linearity and limit of detection were performed per Nanostring protocol. The mRNA expressed below background were filtered from the analysis using cutoffs of mean plus two standard deviations of negative controls and only probes with counts greater than the background threshold were included in the analysis.

### Mass Cytometry

Tissue sections were thawed to room temperature and then baked at 60°C for one hour. The tissue was washed twice in PBS followed by four 5-minute washes in ethanol (50%, 70%, 100% x2), followed by a DEPC water wash. Antigen retrieval was done in 1X Tris EDTA (pH 9.0) at 99°C for 20 minutes using a Hamilton beach steamer. Slides were washed in PBS; tissue was encircled using a PAP pen and allowed to dry for 5 minutes. Tissue was blocked in 3% BSA in PBS for 45 minutes at room temperature in a hydration chamber. Antibody cocktail was prepared in 0.5% BSA in PBS, pipetted onto slide and incubated overnight at 4°C in a hydration chamber. The tissue was then washed twice in 0.2% Triton X-100, followed by two washes in PBS for 8 minutes with slow agitation. The tissue was stained with Intercalator-Ir solution for 30 minutes at room temperature in a hydration chamber. The slides were washed in DEPC water for 5 minutes and air-dried for 20-minutes at room temperature. Tissue slides were stored in a slide box at 4°C until acquired on the CyTOF Hyperion (Standard BioTools).

### Statistical methods

All analyses were performed with GraphPad Prism 9 software. Data was presented as mean ± S.D. or S.E.M. with statistically significant P-values determined by non-parametric Mann-Whitney t-test or Kruskal-Wallis one-way ANOVA as indicated in figure legends.

## Acknowledgements

We would like to thank the members of the Beam Therapeutics Immunology team and the Hatfield Laboratory that contributed to this collective work. Special acknowledgement for J. Decker and the Beam sequencing core, R. Manoukian and the Beam imaging core for their technical expertise. B. Lutz for her expert guidance with spatial transcriptomics at Beam Therapeutics, and BioRender for schematic design. These studies were completed with collaborative research funding from Beam Therapeutics.

## Author Contributions

R.M., C.M., S.H., conceptualized the project and designed the experiments. R.M., S.H., N.N., K.D., M.C., S.C., B.C., A.M., H.S., J.R., F.M., A.C., performed experiments, analyzed results and interpreted data. R.M, C.M., S.H. wrote the manuscript. G.C., provided constructive feedback and secured funding for the project. M.S. provided institutional knowledge and infrastructure at the New England Inflammation and Tissue Protection Institute, provided constructive criticism, and participated in manuscript preparation.

---

**Supplementary Table 1.**
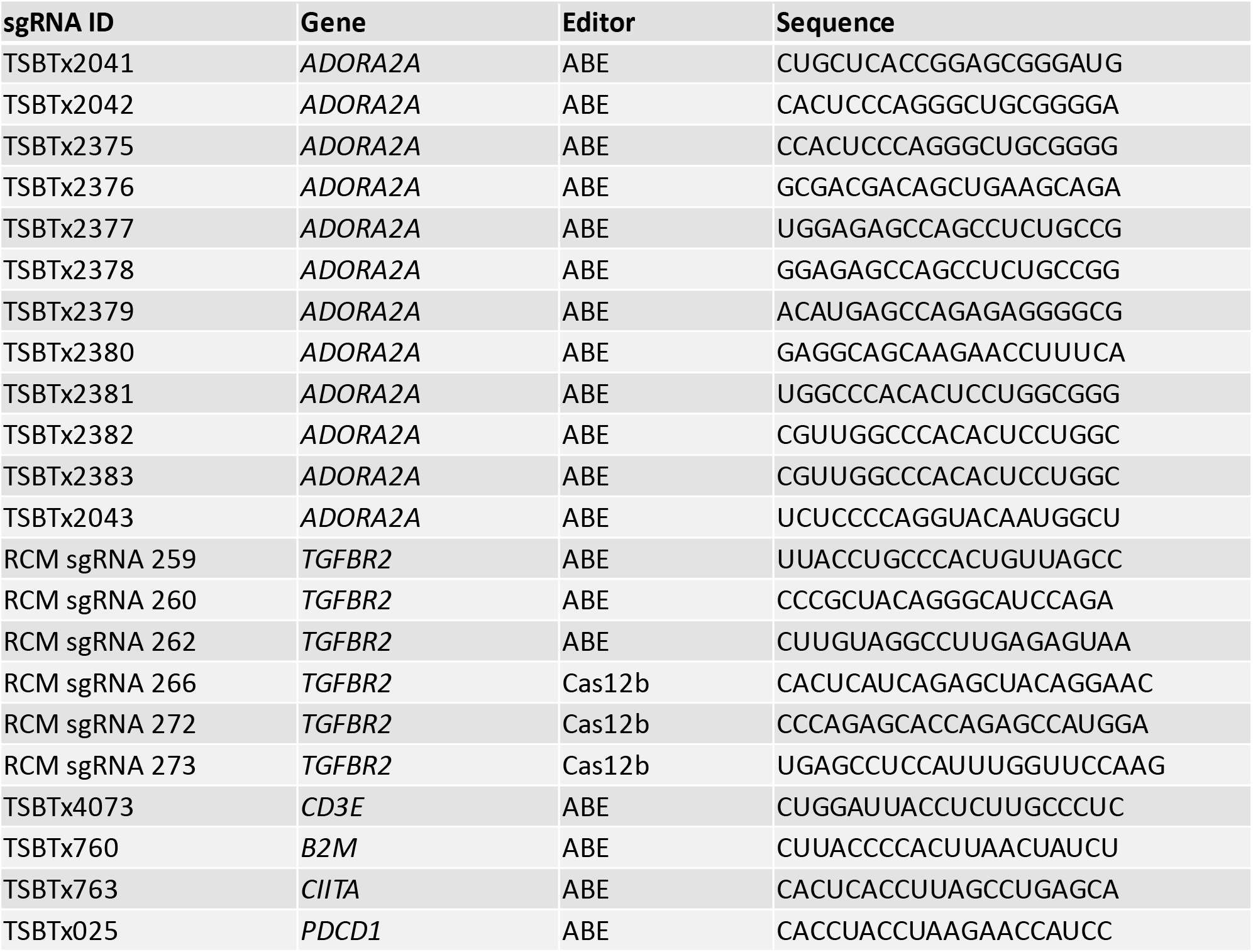

