## Supplementary Figures for "Comprehensive genome editing confers ‘off-the-shelf’ CAR-T cells superior efficacy against solid tumors"

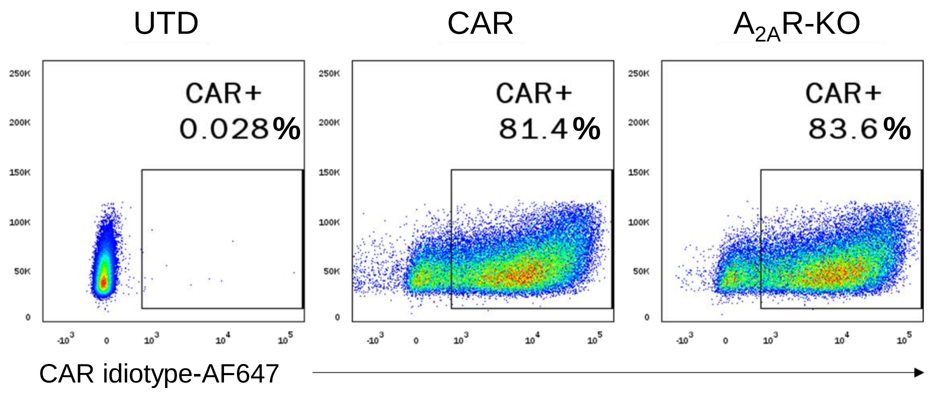


**Supplementary Fig. 1: Genetic ablation of *ADORA2A* does not impact anti-EGFR CAR expression in primary human T cells. a,** Representative flow cytometry plots depicting CAR expression of untransduced (UTD) T cells, unedited (CAR) and $A_{2A}R$-KO CAR-T cells detected with anti-CAR idiotype antibody.

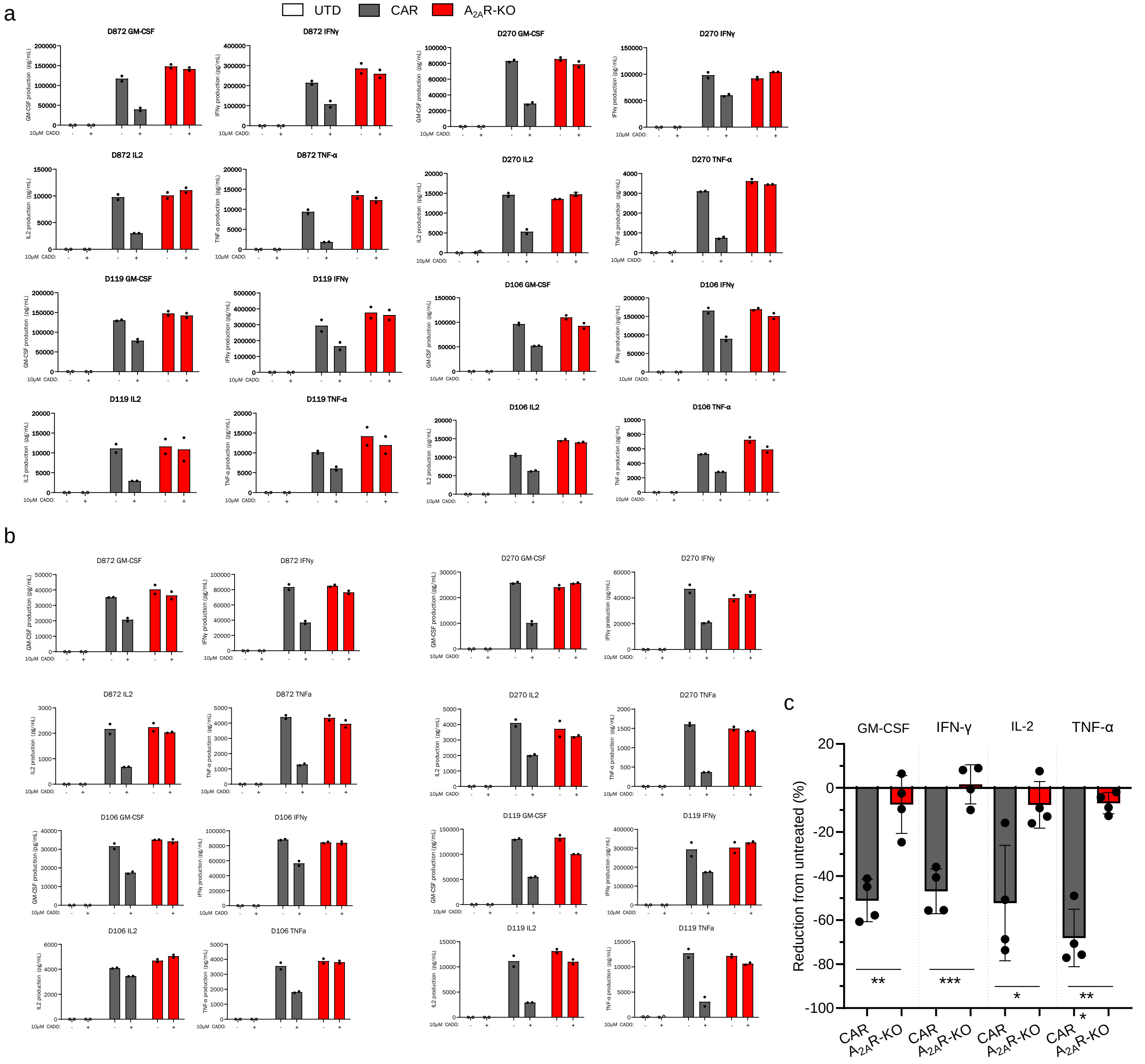


**Supplementary Fig. 2: $A_{2A}$R-KO CAR-T cells are resistant to extracellular adenosine-mediated cytokine suppression *in vitro*. a-c,** GM-CSF, IFN-γ, IL2 and TNF-α production by untransduced (UTD) T cells (white), unedited (gray) or $A_{2A}$R-KO (red) CAR-T cells from four independent biological donors 48 hours after stimulation with $EGFR^{+}$ H226 (**a**) or A549 (**b, c**) tumor cells in the presence (+) or absence (-) of 2-chloroadenosine (cADO) as measured by ELISA. **c,** Magnitude of GM-CSF, IFN-γ, IL2, TNF-α secretion by unedited (gray) and $A_{2A}$R-KO (red) CAR-T cells in the presence of cADO normalized to respective untreated CAR-T cells. For all data, bars represent mean and symbols reflect technical replicates, except **c,** where symbols represent individual donors ± SD. **c**, Mann-Whitney t-test performed to calculate statistical significance, ****P*<0.001, ***P*<0.01, **P*<0.05.

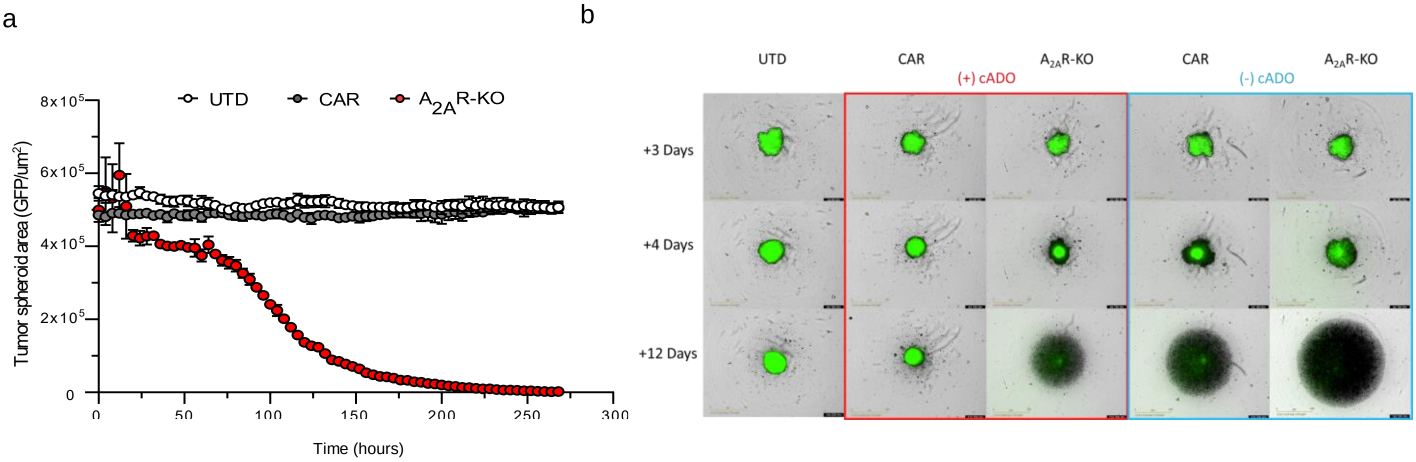


**Supplementary Fig. 3: $A_{2A}$R-KO CAR-T cells retain cytotoxic capacity in the presence of extracellular adenosine *in vitro*. a,** Incucyte-based GFP$^{+}$ H226 tumor spheroid cytotoxicity assay. Untransduced (UTD) T cells (white), unedited CAR0T cells (CAR, gray) or $A_{2A}$R-KO CAR-T cells (red) were cultured with tumor spheroids in the presence of cADO from an independent biological donor. Cytotoxicity calculated as reduction of tumor GFP$^{+}$ area over time. **b,** Representative Incucyte images from (**a**) with cytotoxicity shown 3-, 4-, and 12-days post CAR-T addition in the presence (red) or absence (blue) of cADO. For all data, symbols and error bars reflect mean ± SD of technical replicates.

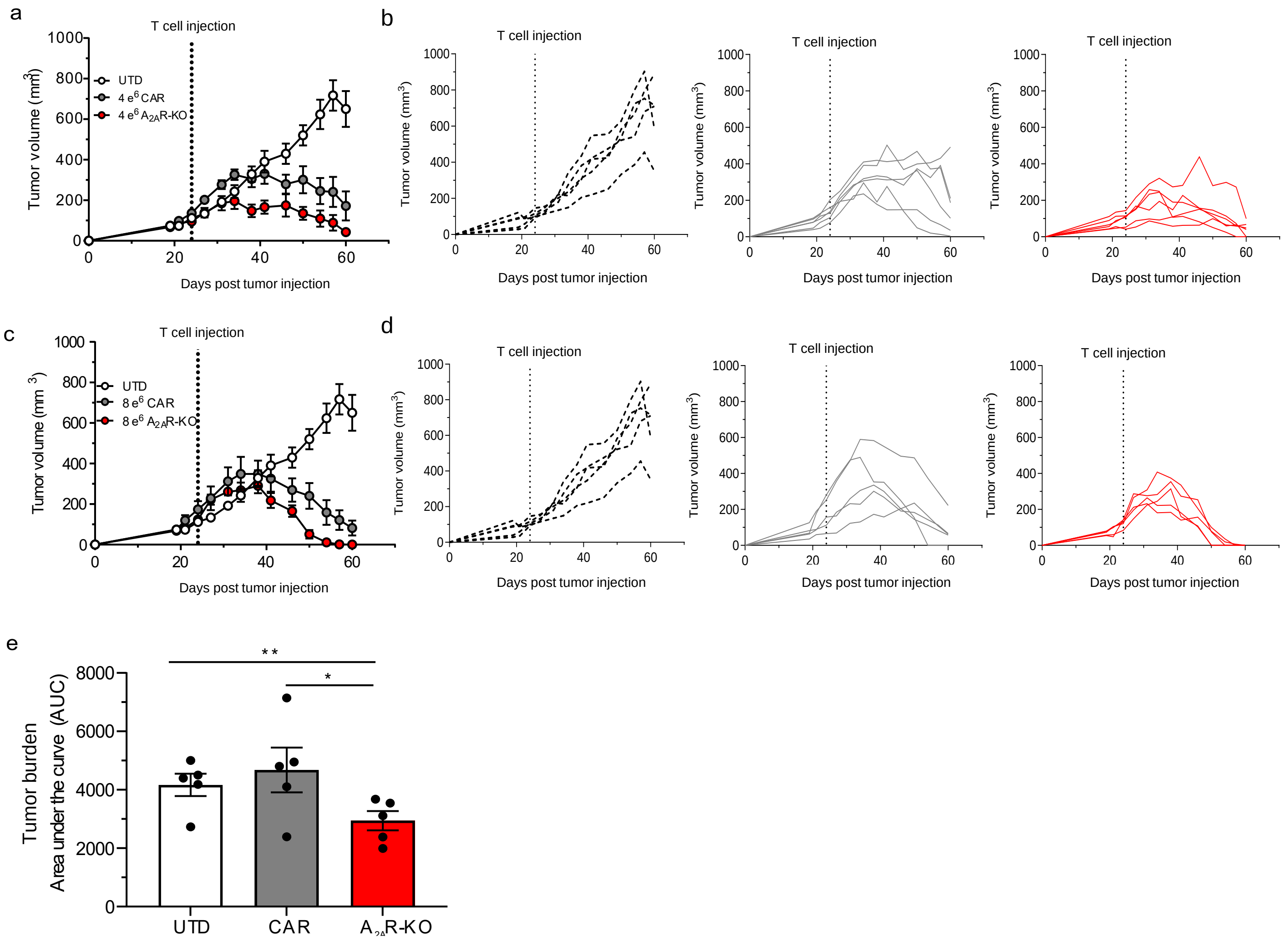


**Supplementary Fig. 4: $A_{2A}$R-KO CAR-T cells demonstrate superior *in vivo* dose-dependent tumor killing kinetics. a-d,** H226 tumor-bearing NCG mice were infused with $4x10^6$ (**a,b**) or $8x10^6$ (**c,d**) untransduced (UTD) T cells (white, $n$ = 5), unedited CAR-T cells (gray, $n$ = 6) and $A_{2A}$R-KO CAR-T (red, $n$ = 6) when tumors reached an average volume of $150mm^3$. Longitudinal group averages (**a,c**) and individual mouse (**b,d**) tumor volume was measured via calipers. **e,** Cumulative tumor burden was determined by calculating area under the curve from H226 tumor-bearing NCG mice that were treated with $2x10^6$ UTD T cells, unedited CAR-T cells, or $A_{2A}$R-KO CAR-T cells. **b,d,e** lines and symbols represent individual mice, **a,c,** symbols represent group mean ± S.E.M. **e**, Kruskal-Wallis test performed to calculate statistical significance, **$P$<0.01, *$P$<0.05.

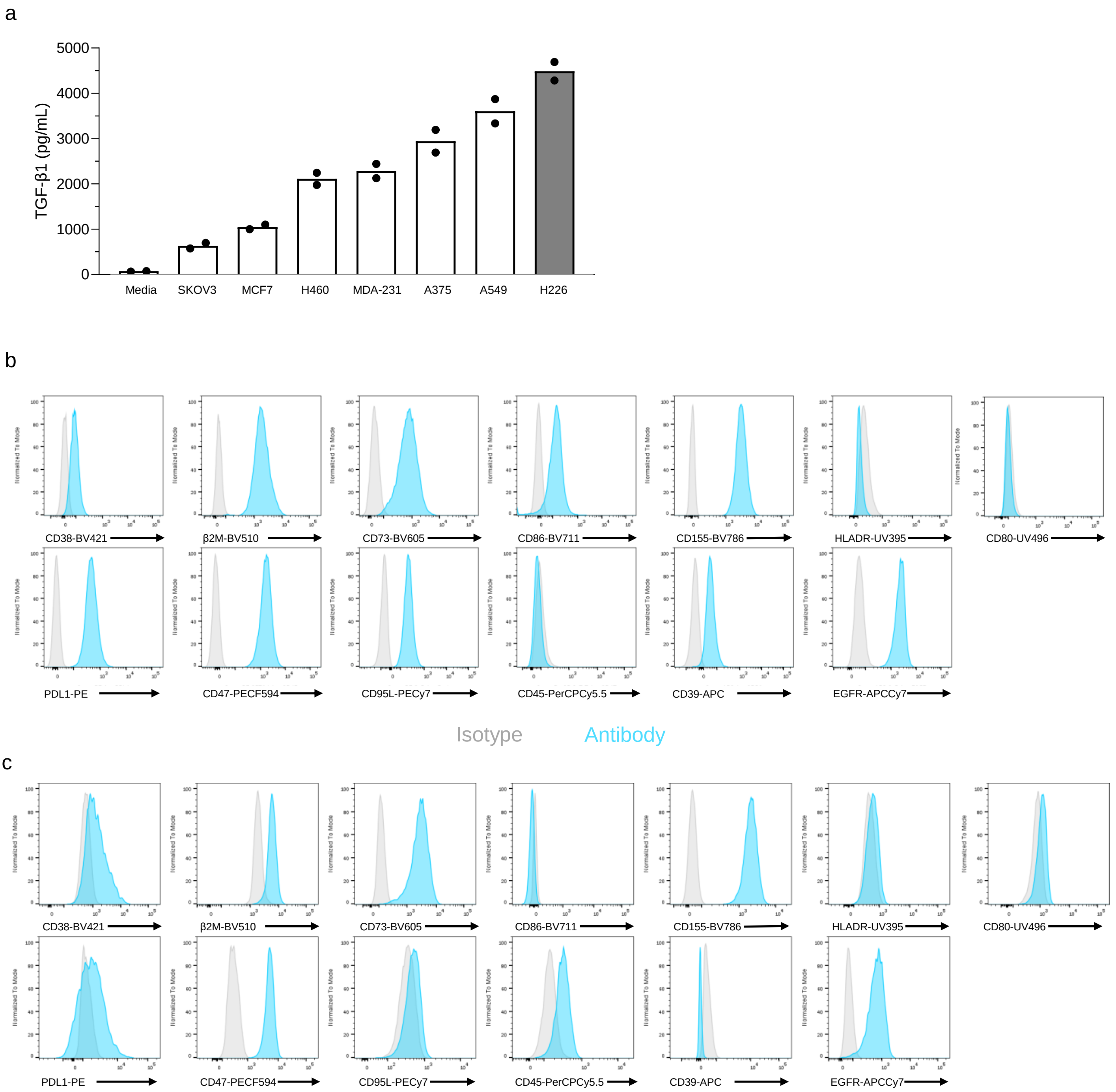


**Supplementary Fig. 5: *In vitro* tumor phenotyping unveils expression of alternative immunosuppressive pathways. a,** TGF-β1 production by the indicated solid tumor cancer cell lines detected via ELISA after 24 hours in culture. Bars reflect mean and symbols represent technical replicates. **b,c,** Flow cytometry histograms depicting expression of cell surface proteins on H226 (**b**) or A549 (**c**) representing major potential immunosuppressive pathways. Isotype (gray), specific antibody (blue).

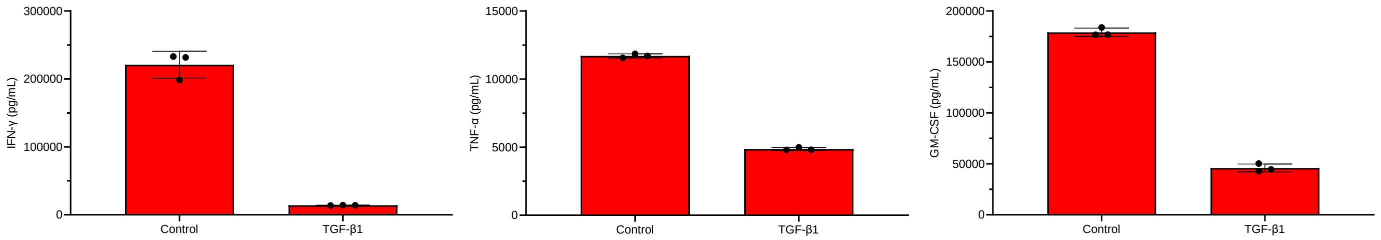


**Supplementary Fig. 6: $A_{2A}$R-KO CAR-T cell pro-inflammatory cytokine release is suppressed by TGF-β.** IFN-γ, TNF-α and GM-CSF cytokine production by $A_{2A}$R-KO CAR-T cells 48 hours post stimulation with H226 tumor cells in the presence (TGF-β1) or absence (control, DMSO) of exogenous TGF-β. Bars represent mean, and symbols reflect technical replicates ± SD.

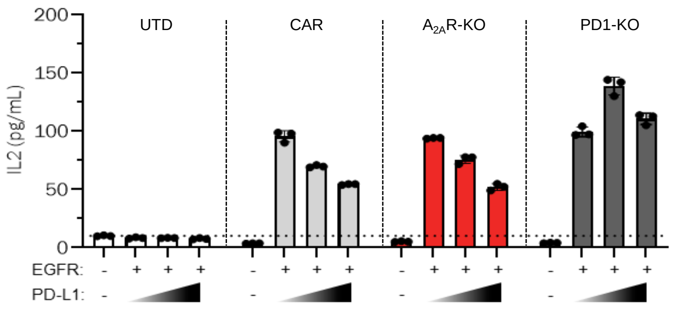


**Supplementary Fig. 7: Genetic deletion of PD-1 protects CAR-T cells from PDL1-mediated inhibition.** IL2 production by untransduced (UTD) T cells, or unedited (light gray), $A_{2A}$R-KO (red) and PD1-KO (dark gray) CAR-T cells 48 hours post-stimulation with recombinant human EGFR and PD-L1 conjugated beads from an additional independent biological T cell donor measured by ELISA. Bars represent mean, symbols reflect technical replicates and error bars are ± SD.

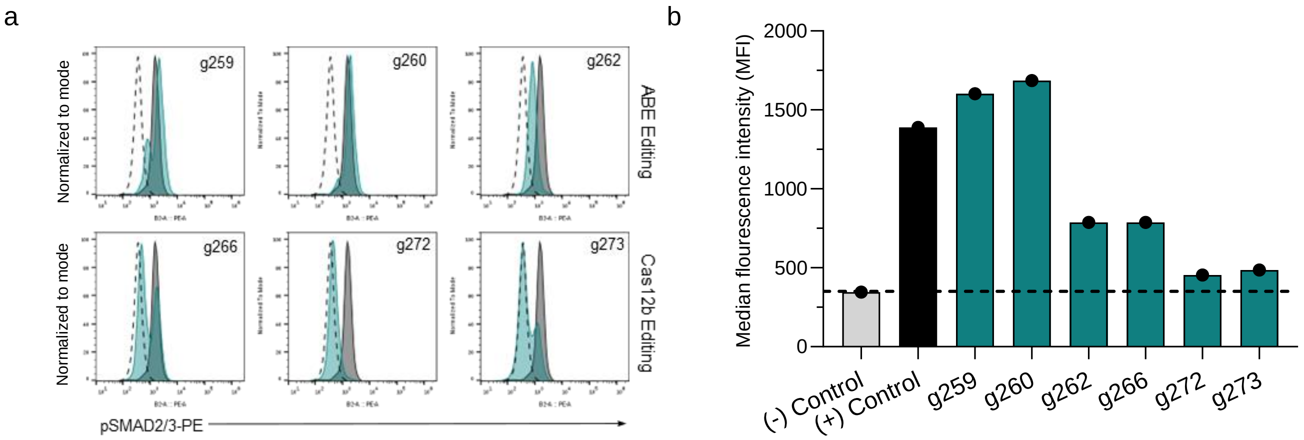


**Supplementary Fig. 8: *TGFBR2* sgRNA screen confirms Cas12b-mediated ablation of downstream TGFβR signaling. a-b,** Control CAR-T cell were unedited (black), and edited CAR-T cells were electroporated with *TGFBR2*-specific single guide RNAs (green) complexed with mRNA encoding either ABE or Cas12b. CAR-T cells were then treated with exogenous TGF-β compared to unedited CAR-T cells rerated with DMSO (dashed line). Histograms depict expression levels of phosphorylated SMAD2/3 (pSMAD2/3) (**a**) and median fluorescent intensity of pSMAD2/3 (**b**) where (-) Control denotes unedited CAR-T cells treated with DMSO and (+) Control denotes unedited CAR-T cells treated with TGF-β. g259-g273 represent individual sgRNAs. Bars represent mean and symbols reflect technical replicates.

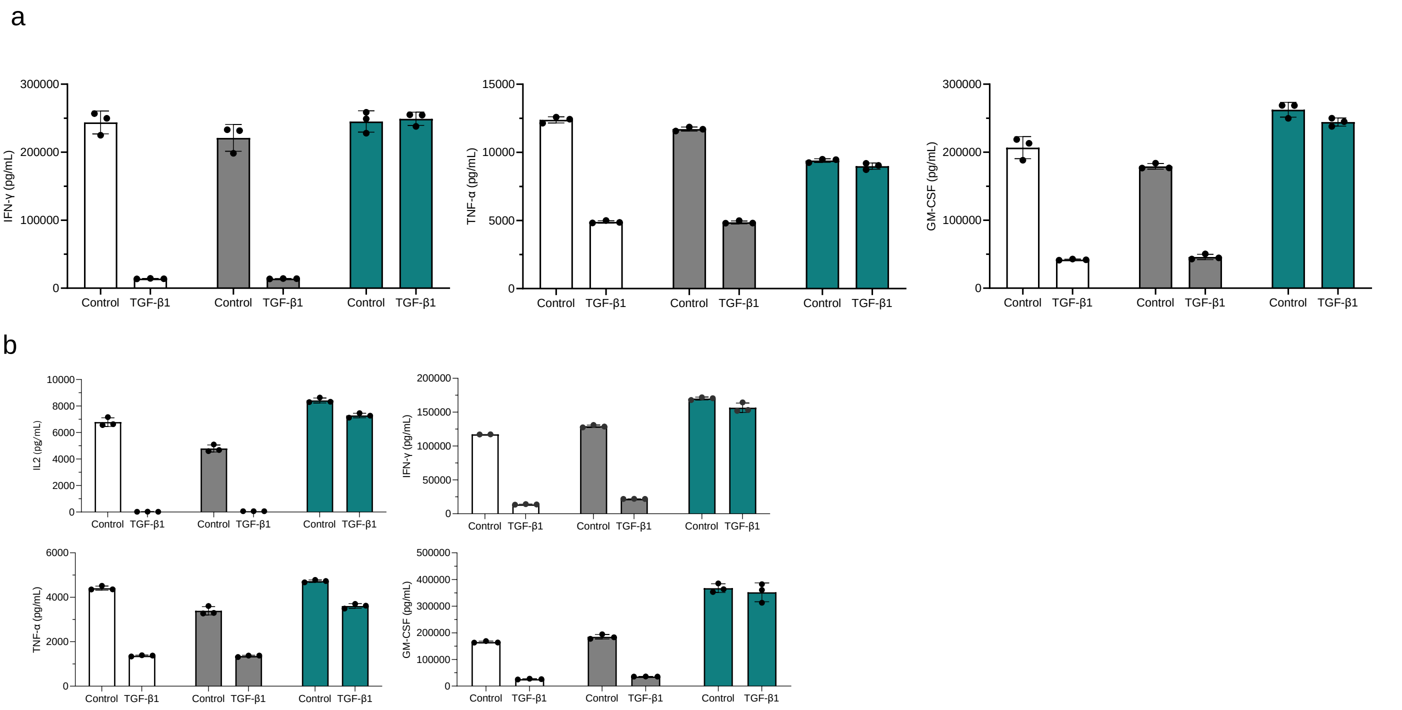


**Supplementary Fig. 9: Genetic ablation of TGFβRII rescues CAR-T cells from TGFβ-mediated immunosuppression. a,** IL2, IFN-γ, TNF-α and GM-CSF cytokine production by unedited (white), $A_{2A}$R-KO (gray) and TGFβRII-KO (green) CAR-T cells 48 hours post stimulation with H226 tumor cells in the presence of exogenous TGF-β1 or DMSO (control). **b,** same as in (**a**) except performed with an additional independent T cell donor. For all data, bars represent mean, symbols reflect technical replicates and error bars ± SD.

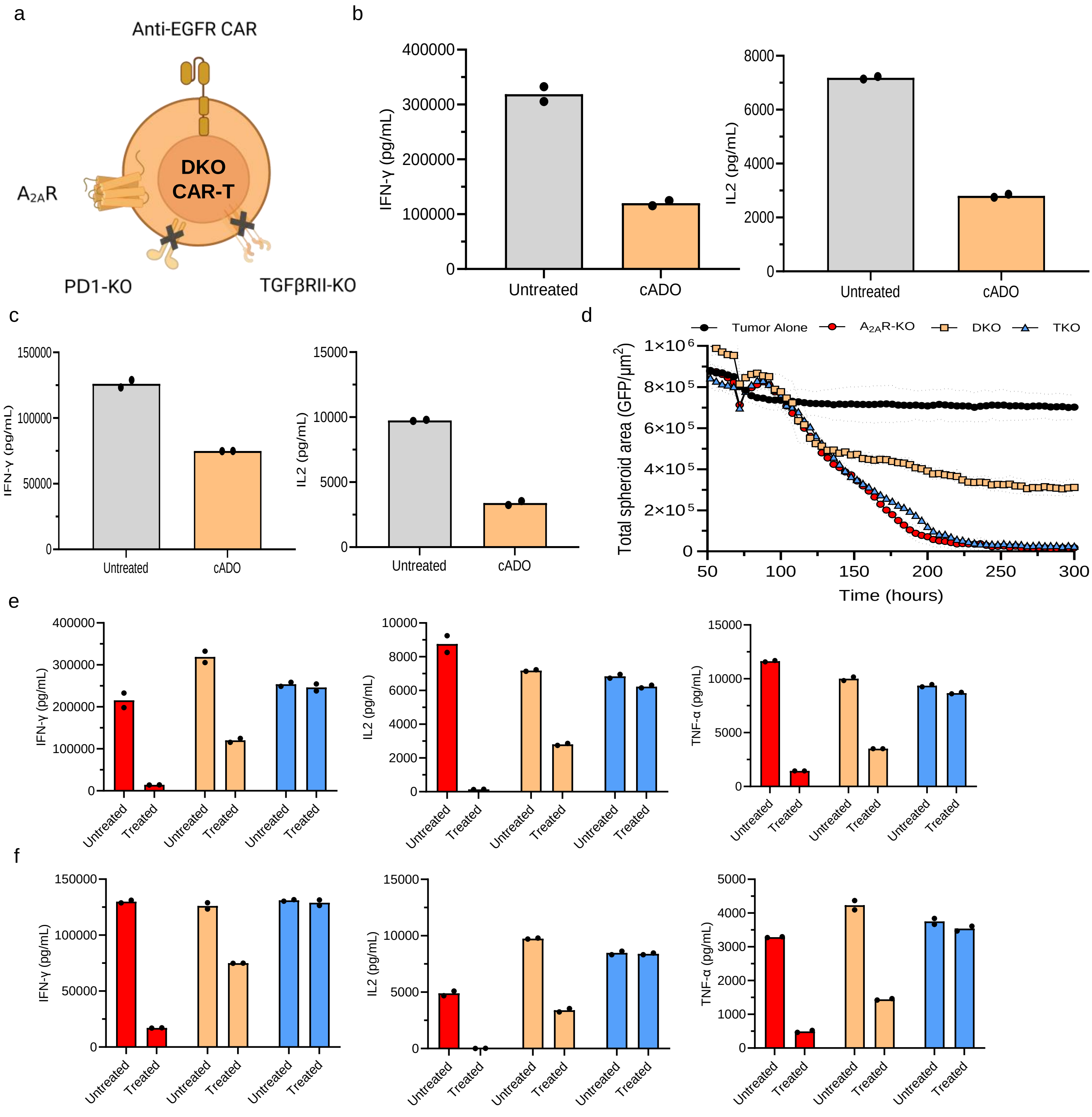


**Supplementary Fig. 10: PD-1 and TGFβR2 double knockout CAR-T cells are potently suppressed by biochemical negative regulators. a,** Schematic of *PDCD1* and *TGFBR2* double knock out (DKO) CAR-T cells depicting loss of PD-1 and TGFβRII protein expression. **b,** IFN-γ and IL2 production by DKO CAR-T cells 48 hours post stimulation with H226 tumor cells in the presence of adenosine (cADO, orange) compared to DMSO-treated control (untreated, gray) measured by ELISA. **c,** Cytokine production as in (**b**) from an independent T cell donor. **d,** Incucyte live imaging GFP$^+$ H226 tumor spheroid cytotoxicity of $A_{2A}R$-KO (red circle), DKO (orange square), and TKO (blue triangle) CAR-T cells compared to tumor alone (black circle) in the presence of cADO. Cytotoxicity was measured as a decrease in cumulative tumor GFP$^+$ area over time. **e,** IFN-γ, IL2 and TNF-α production by $A_{2A}R$-KO (white), DKO (orange) and TKO (gray) CAR-T cells 48 hours post stimulation with H226 in the presence of DMSO (control) or triple suppression comprising cADO and exogenous PDL-1 and TGF-β (treated). **f,** Cytokine production as in (**e**) from an independent T cell donor. For all data, bars represent mean, symbols indicated technical replicates and erro bars are ± SD.

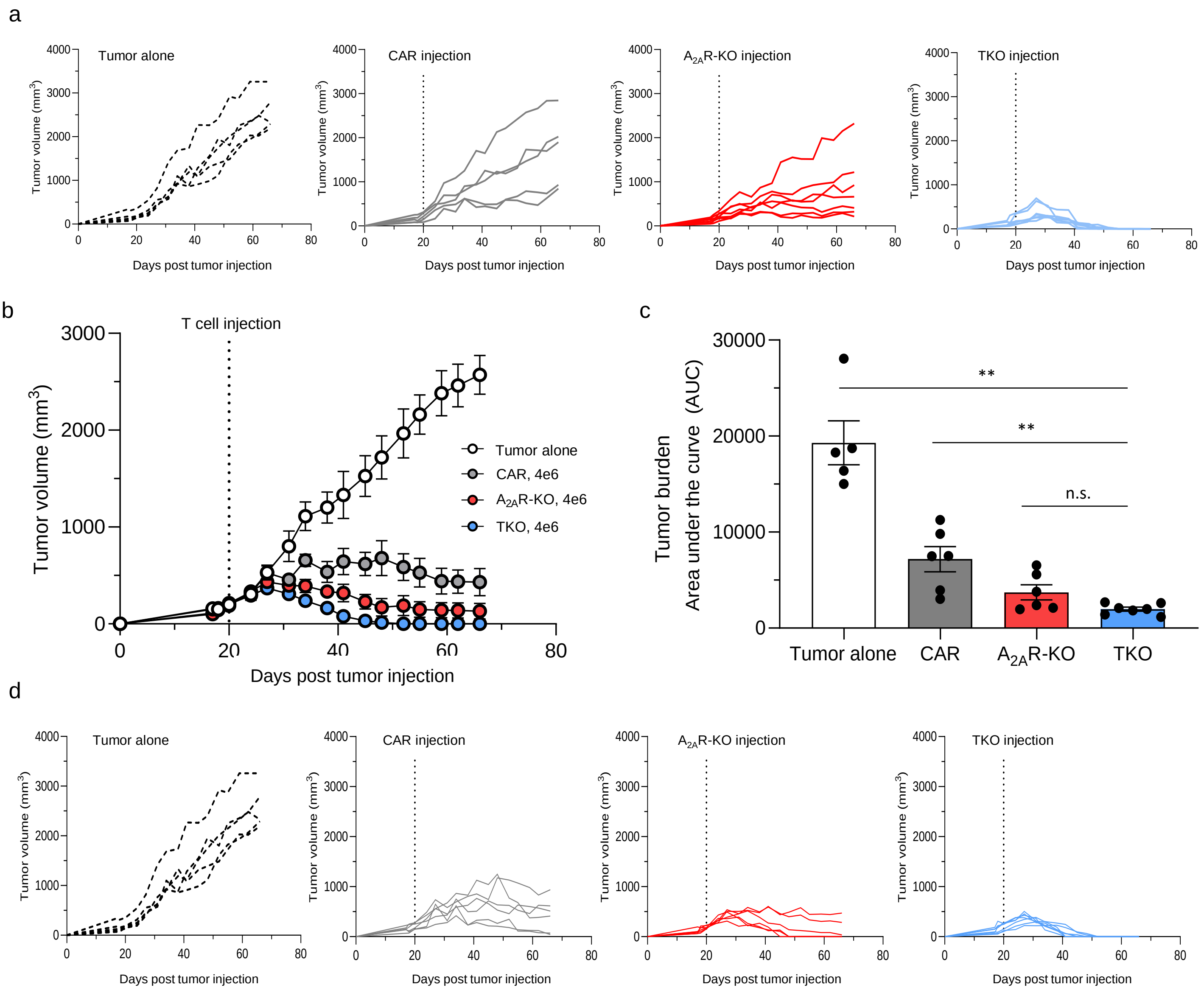


**Supplementary Fig. 11: TKO CAR-T cells exhibit increased anti-tumor activity in NCG mice. a,** Tumor volume of group individual mice from $2x10^6$ cell dose of untreated (white) or unedited (gray), $A_{2A}R$-KO (red) and TKO (blue) CAR-T cells. **b,** $4x10^6$ dose of untreated (white, $n$ = 5) or unedited CAR (gray, $n$ = 6) and $A_{2A}R$-KO (red, $n$ = 6) and TKO (blue, $n$ = 7) CAR-T cells infused via tail vein into H226-bearing NCG mice when tumor volume reached an average of $150mm^3$. **c,** Reduction of cumulative tumor burden determined by area under the curve from $4x10^6$ CAR-T dose. **d,** Tumor burden from individual mice from (**b**). **e,** Representative flow cytometry plots of $CD2^+ZsGreen^+$ CAR, $A_{2A}R$-KO and TKO CAR-T cells detected in H226 tumors resected from NCG mice 7 days post CAR-T infusion. For all data, symbols and error bars reflect individual mice ± SD, except **b**,**c,** here symbols and bars represent group mean ± S.E.M. **c**, Mann-Whitney test performed to calculate statistical significance, ***P*<0.01, **P*<0.05.

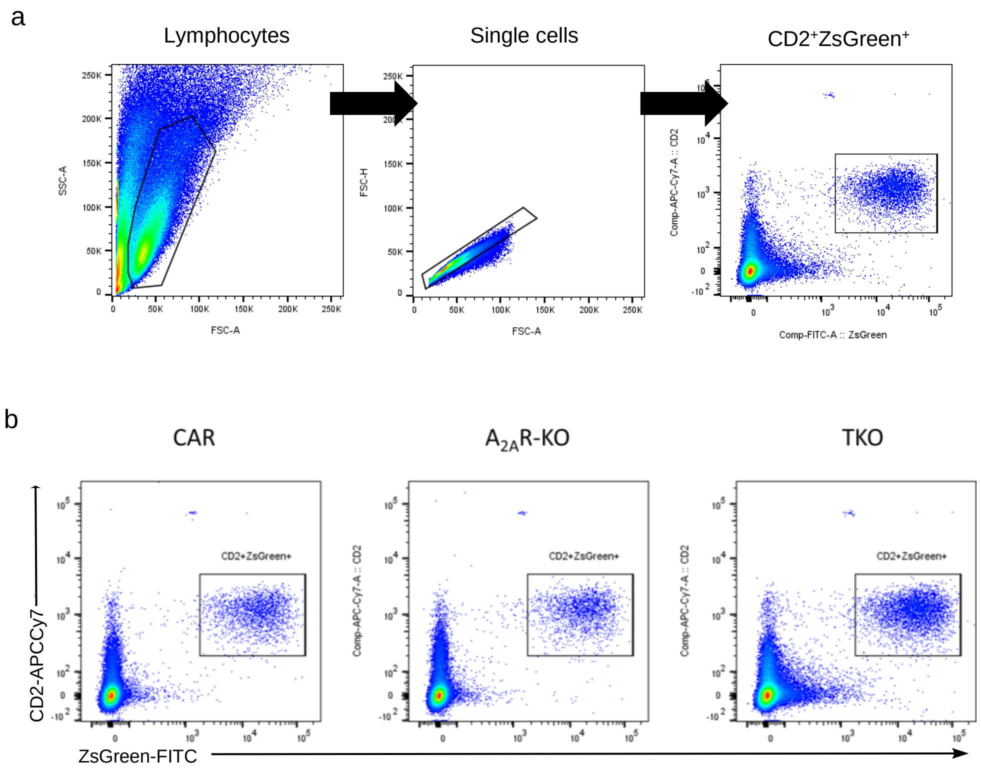


**Supplementary Fig. 12: CAR-T cells are detected in H226 tumors resected from NCG mice. a,** Gating strategy used to identify ZsGreen$^+$ CAR-T cells isolated from H226 tumors resected from NCG mice 7 days post CAR-T infusion.. **b,** Representative flow cytometry plots of CD2$^+$ZsGreen$^+$ CAR-T cells, $A_{2A}$R-KO CAR-T cells and TKO CAR-T cells.

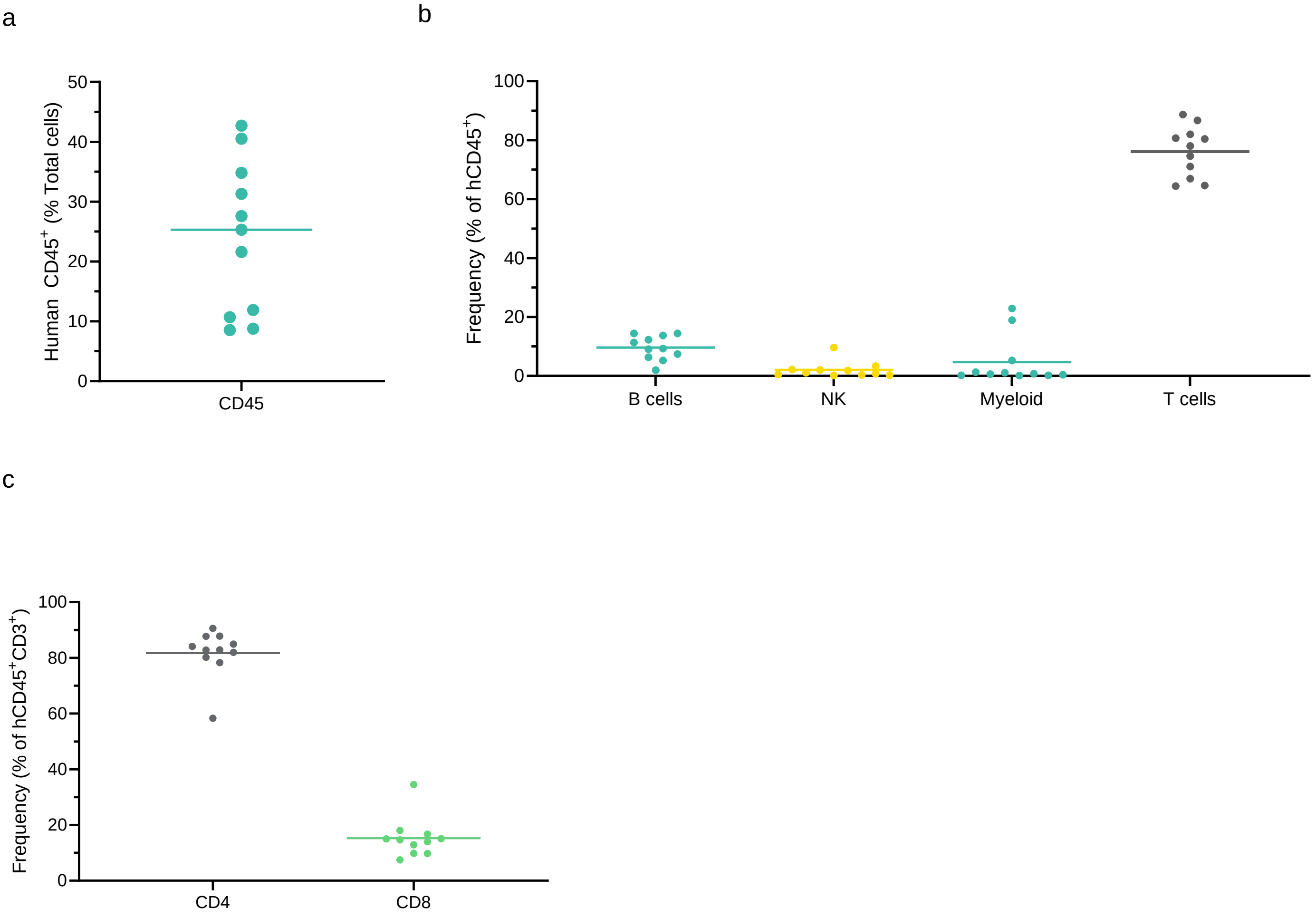


**Supplementary Fig. 13: Characterization of the human immune compartment in humanized-NCG mice. a-c,** Flow cytometry analysis of humanized NCG (huNCG) mouse blood identifying human CD45$^+$ cells (**a**), human immune cell subsets including CD19$^+$ B cells, CD56$^+$ NK cells, CD33$^+$ myeloid cells and CD3$^+$ T cells (**b**) with CD4$^+$ and CD8$^+$ expression T cell subset phenotyping (**c**) prior to CAR-T cell infusion. For all data, symbols represent individual mice, and bar represents mean.

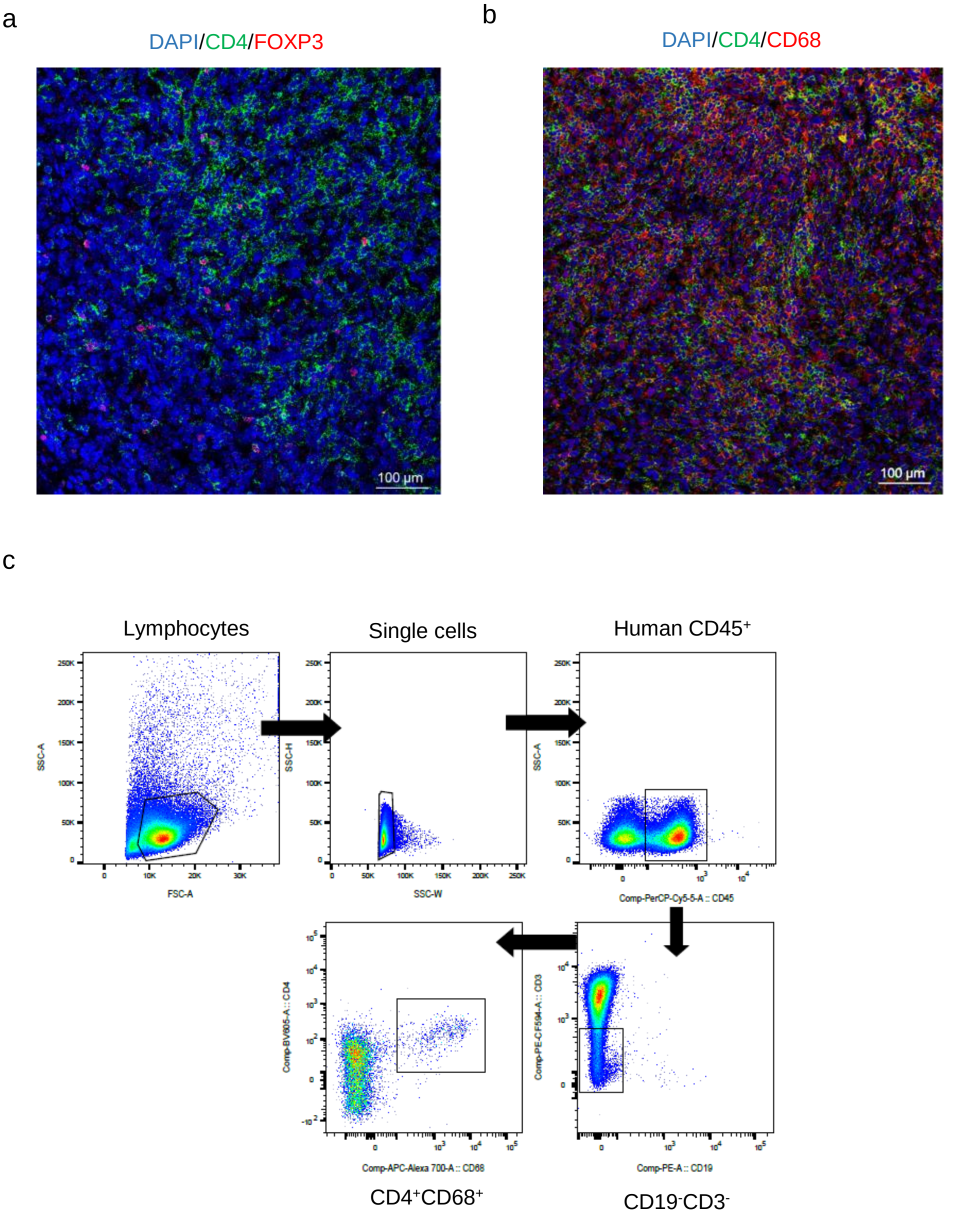


**Supplementary Fig. 14: Human immune cells detected in the TME of H226-bearing humanized-NCG mice. a,b,** Mass cytometry of H226 tumors resected from huNCG mice identifying $CD4^+$ (green) $FoxP3^+$ (red) cells in (**a**) and $CD68^+$ (red) cells (**b**) present in the TME. Nucleated cell (DAPI, blue). **c,** Representative flow cytometry gating strategy used to identify $CD45^+CD3^-CD19^-CD4^+CD68^+$ cells isolated from H226 tumors resected from huNCG mice.

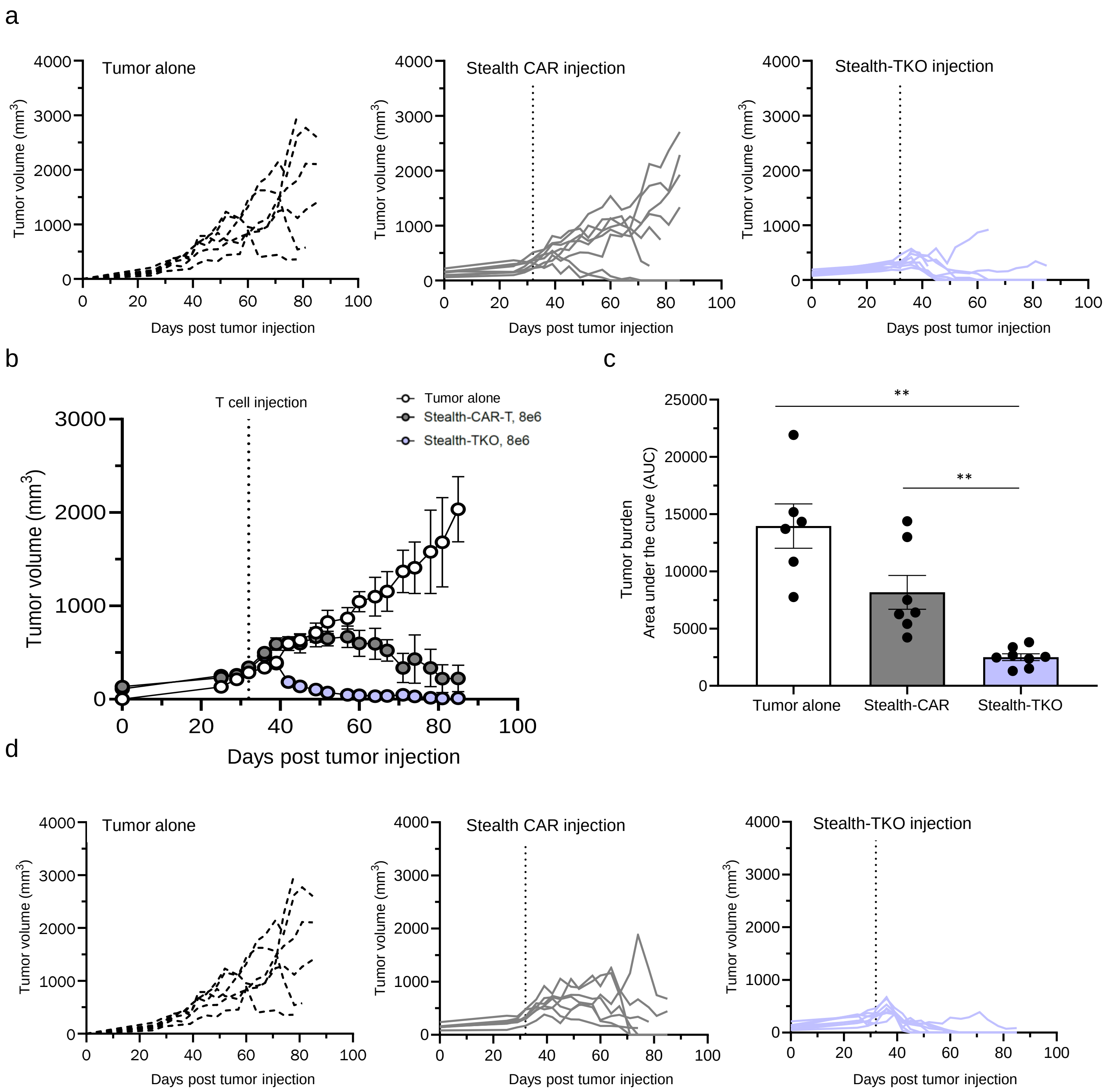


**Supplementary Fig. 15: Stealth-TKO CAR-T cells demonstrate dose-dependent anti-tumor control in humanized mice. a,** Tumor volume of individual mice from 4x10$^6$ cell dose of Stealth-CAR (gray), and Stealth-TKO (purple) CAR-T cells in humanized NCG (huNCG) mice compared to an untreated control group (dashed). **b,** 8x10$^6$ dose of Stealth-CAR (gray, *n* = 7) and Stealth-TKO (purple, *n* = 8) CAR-T cells infused via tail vein into H226-bearing NCG mice when tumor volume reached an average of 150mm$^3$ compared to untreated (control) mice. **c,** Reduction of cumulative tumor burden determined by area under the curve from 8x10$^6$ CAR-T dose. **d,** Tumor burden from individual mice from (**b**)**.** For all data, symbols and error bars reflect individual mice ± SD, except **b,c**, where symbols and error bars represent mean ± S.E.M. **c,** Mann-Whitney test used to calculate statistical significance, **$P$<0.001.
